# The role of the anterior intraparietal sulcus and the lateral occipital cortex in fingertip force scaling and weight perception during object lifting

**DOI:** 10.1101/2019.12.20.883918

**Authors:** Vonne van Polanen, Guy Rens, Marco Davare

## Abstract

Skillful object lifting relies on scaling fingertip forces according to the object’s weight. When no visual cues about weight are available, force planning relies on recent lifting experience. Recently, we showed that previously lifted objects also affect weight estimation, as objects are perceived to be lighter when lifted after heavy objects compared to light ones. Here, we investigated the underlying neural mechanisms mediating these effects. We asked participants to lift objects and estimate their weight. Simultaneously, we applied transcranial magnetic stimulation (TMS) during the dynamic loading or static holding phase. Two subject groups received TMS of either the anterior intraparietal sulcus (aIPS) or lateral occipital area (LO), known to be important nodes in object grasping and perception. We hypothesized that TMS-induced disruption of aIPS and LO would alter force scaling and weight perception. Contrary to our hypothesis, we did not find effects of aIPS or LO stimulation on force planning or weight estimation caused by previous lifting experience. However, we found that TMS of both areas increased grip forces, but only when applied during dynamic loading, and decreased weight estimation, but only when applied during static holding, suggesting time-specific effects. Interestingly, our results also indicate that TMS over LO, but not aIPS, affected load force scaling specifically for heavy objects, which further indicates that planning of load and grip forces might be controlled differently. These findings provide new insights on the interactions between brain networks mediating action and perception during object manipulation.

**NEW & NOTEWORTHY:** This article provides new insights into the neural mechanisms underlying object lifting and perception. Using transcranial magnetic stimulation during object lifting, we show that effects of previous experience on force scaling and weight perception are not mediated by the anterior intraparietal sulcus nor the lateral occipital cortex (LO). In contrast, we highlight a unique role for LO in load force scaling, suggesting different brain processes for grip and load force scaling in object manipulation.

## 1 INTRODUCTION

Every day we skillfully manipulate various objects using our hands. In order to lift an object skillfully, one has to precisely adjust fingertip forces to the weight of the object. That is, sufficient grip force, i.e. the force perpendicular to the object surface, has to be applied to avoid slipping of the object. In addition, the load force, i.e. the vertical force, has to overcome gravity and be equal to object weight in static object holding. Object weight can be predicted from object properties, such as size or material, to form a motor plan and ensure a smooth lifting motion. Because feedback processes are slow, anticipatory scaling of fingertip forces results in more fluent lifts compared to feedback-driven movements. If the motor plan is incorrect, feedback processes can be used to quickly adapt the motor plan and apply the correct forces (Johansson and Westling 1988).

When the weight of an object cannot be inferred from object properties, one usually relies on previous lifting experience with that object, which is often referred to as sensorimotor memory (Johansson and Westling 1988). For instance, if a heavy object has been lifted, the next lift on an object with identical appearance will be scaled towards the heavy weight as well. The sensorimotor memory can be maintained for hours (Flanagan et al. 2001; Green et al. 2010; Nowak et al. 2007), transferred between hands (Chang et al. 2008; Gordon et al. 1994; Nowak et al. 2005) and has a neural representation in the primary motor cortex (Chouinard et al. 2005; Loh et al. 2010).

The actual weight of an object can only be unequivocally determined after it has been lifted from the table. Recently, it has been suggested that sensorimotor memory effects could be linked to weight estimation. When an object was lifted after a heavy object, it was judged to be lighter than when lifted after a light one (van Polanen and Davare 2015b). Interestingly, force scaling parameters correlated with this perceptual bias and both effects of force scaling and weight perception increased with longer sequences of lifts (i.e. increasing the magnitude of sensorimotor memory effects), suggesting that the correction to fingertip forces and weight perception are associated. This perceptual bias depending on previous object weight has been replicated and was shown to transfer across hands (Maiello et al. 2018). Furthermore, a similar effect was shown for torque planning, which also affected heaviness and weight distribution estimation when lifting objects with an unequal weight distribution (Schneider et al. 2019). All in all, these studies suggest a link between force scaling and perception of object weight.

When acquiring sensory information about object weight, the dynamic phase of movements might be especially important. Since corrections to force planning based on sensory feedback mainly take place during the loading phase, i.e. between object contact and lift-off (Johansson and Flanagan 2009), it is possible that this phase is critical for building up sensorimotor memory and the formation of a weight estimate. Indeed, it has been shown that sensorimotor memory of force scaling is based on information acquired during the lifting phase, not the holding phase (van Polanen and Davare 2019). In addition, when observing lifting movements of others, the lifting phase was found to be important for making judgements of object weight (Hamilton et al. 2007). Finally, the relation between action planning and weight perception (Schneider et al. 2019; van Polanen and Davare 2015b) suggests that this phase could also be important for weight perception during the execution of object lifting. More specifically, van Polanen and Davare (2015b) showed that the perceptual bias was absent if objects were not lifted but a weight was passively pressed on the hand.

The neural pathways processing vision for action and perception have classically been divided into a dorsal and ventral stream. More specifically, the dual stream theory proposes that visual information used for spatially locating the object and planning the action is processed in the dorsal stream, running from visual cortex to parietal areas, whereas information for object recognition is managed by the ventral stream, running from visual cortex towards the temporal cortex (Goodale and Milner 1992; Milner and Goodale 2008; Ungerleider and Mishkin 1982). A similar division between action-perception processing has been suggested for somatosensory perception (Dijkerman and de Haan 2007). It has also been argued that both visual streams do interact heavily (Cloutman 2013; Schenk and McIntosh 2010), especially as motor skill demands increase (van Polanen and Davare 2015a).

In the present study, we wanted to investigate the interplay between the dorsal and ventral stream during the execution of an action-perception task. We focused on two key areas in the dorsal and ventral stream. First, we hypothesized that the anterior intraparietal sulcus (aIPS), part of the dorsal stream, could be an important node for controlling forces during object lifting and the correction of erroneously scaled forces. Indeed, this area is known to be involved in grip force scaling (Dafotakis et al. 2008; Davare et al. 2007). Regarding object perception, a similar status might be allocated to the lateral occipital (LO) cortex, which is part of the ventral stream. We hypothesized that LO could mediate weight estimation, since this area is important for object recognition in both the visual and haptic modality (Amedi et al. 2002; Amedi et al. 2001) and is involved in the representation of object weight (Gallivan et al. 2014).

Furthermore, it is known that the temporal and parietal cortices are connected, based on primate (Borra et al. 2008; Borra et al. 2010; Distler et al. 1993) and human studies (Budisavljevic et al. 2018; Ramayya et al. 2010) which could provide the basis for communication between dorsal and ventral streams. In addition, aIPS has been suggested to play an intermediate role in somatosensory action-perception interactions (Sedda and Scarpina 2012). Therefore, we hypothesized that aIPS and LO could be important nodes in the link between action and perception processes.

The aim of the current study was to investigate the role of aIPS and LO in force scaling and weight perception. Specifically, we wanted to study how these areas are involved in the relation between action planning and perception from previous experience, where force scaling correlated with weight estimation (van Polanen and Davare 2015b). To do this, participants performed an object lifting task and were asked to estimate object weight. To alter their force planning, we varied the order in which light and heavy objects were lifted. Transcranial magnetic stimulation (TMS) during the dynamic loading phase or static holding phase was used to disrupt aIPS and LO and infer their causal role. We expected that disruption induced by TMS of aIPS might not only affect fingertip force scaling (Dafotakis et al. 2008; Davare et al. 2007), but also affect weight perception through connections with LO. When corrections need to be applied to the planned fingertip forces, aIPS might send information to LO which could lead to perceptual weight biases. In return, LO might provide information to aIPS to plan fingertip forces based on known object weight information. More specifically, we expected that aIPS stimulation would reduce both the force corrections and the perceptual bias from previous lifting experience and alter the relation between force scaling and weight perception whereas LO would only affect the perceptual bias. Furthermore, we hypothesized that the involvement of these areas would be more pronounced in the dynamic loading phase compared to the static holding phase.

## 2 METHODS

### 2.1 Participants

30 right-handed participants took part in the study. They were divided into two groups, an aIPS stimulation group (8 males, 7 females, 21±2.2 years) and an LO stimulation group (6 males, 9 females, 21±2.8 years). Their right-handedness was accessed using the Edinburgh handedness questionnaire (Oldfield 1971), which provided a mean L.Q. of 0.9±0.13. All participants gave written informed consent before participation and the study was approved by the local ethical committee of the Biomedical Sciences group at KU Leuven.

### 2.2 TMS procedure

The experiment was divided into two sessions. In the first session, participants were scanned in a 3T MR scanner (Philips Achieva, Philips Healthcare). A high-resolution 3D T1-weighted image was obtained with the following parameters: (TR=9.7 ms, TE=4.6 ms, field of view = 256 × 256 mm^2^, 192 slices, voxel size = 0.98 × 0.98 × 1.2 mm^3^). The images were transferred to Brainsight (Rogue Research), which was used for online neuronavigation throughout the experiment. For the aIPS group, TMS stimulation sites were anatomically determined as the intersection between the postcentral and the intraparietal sulcus (see (Frey et al. 2005), mean MNI coordinates −46±3, −37±5, 46±4). The orientation of the TMS coil was positioned perpendicular to the intraparietal sulcus, with the handle pointing backwards (Figure 2). For the LO group, Talairach coordinates from literature (Amedi et al. 2001) were used, converted to MNI (−47, −61, −16) and located on individual structural MR images. For this group, a similar orientation was used as in the aIPS group, with a slight clockwise rotation, if necessary, to avoid contact of the handle with the shoulder (see Figure 2 for an example).

**Figure 1.**
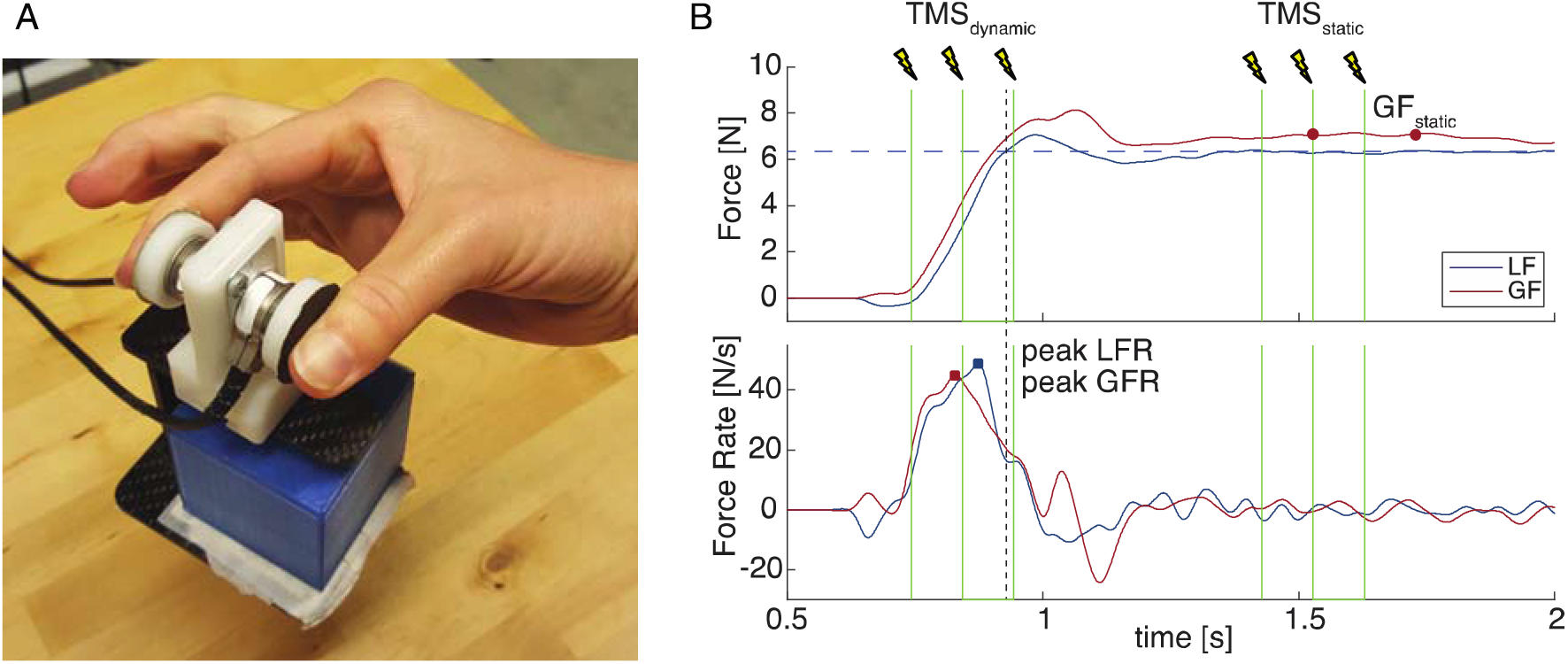
**A.** Manipulandum with force sensors. Different objects can be placed in the basket to change object weight. **B.** Example of trial with force traces of grip and load force (GF, LF; top panel) and force rates (bottom panel). TMS was applied as a burst of 3 pulses at 10Hz, at contact (TMS_dynamic_) or 500 ms after lift-off (TMS_static_). Force parameters were calculated as peak force rates (peak LFR, peak GFR) and grip force during static holding (GF_static_; average between 600-800 ms after lift-off, red circles). Dashed horizontal line indicates lift-off, dashed vertical line indicates object weight.

**Figure 2.**
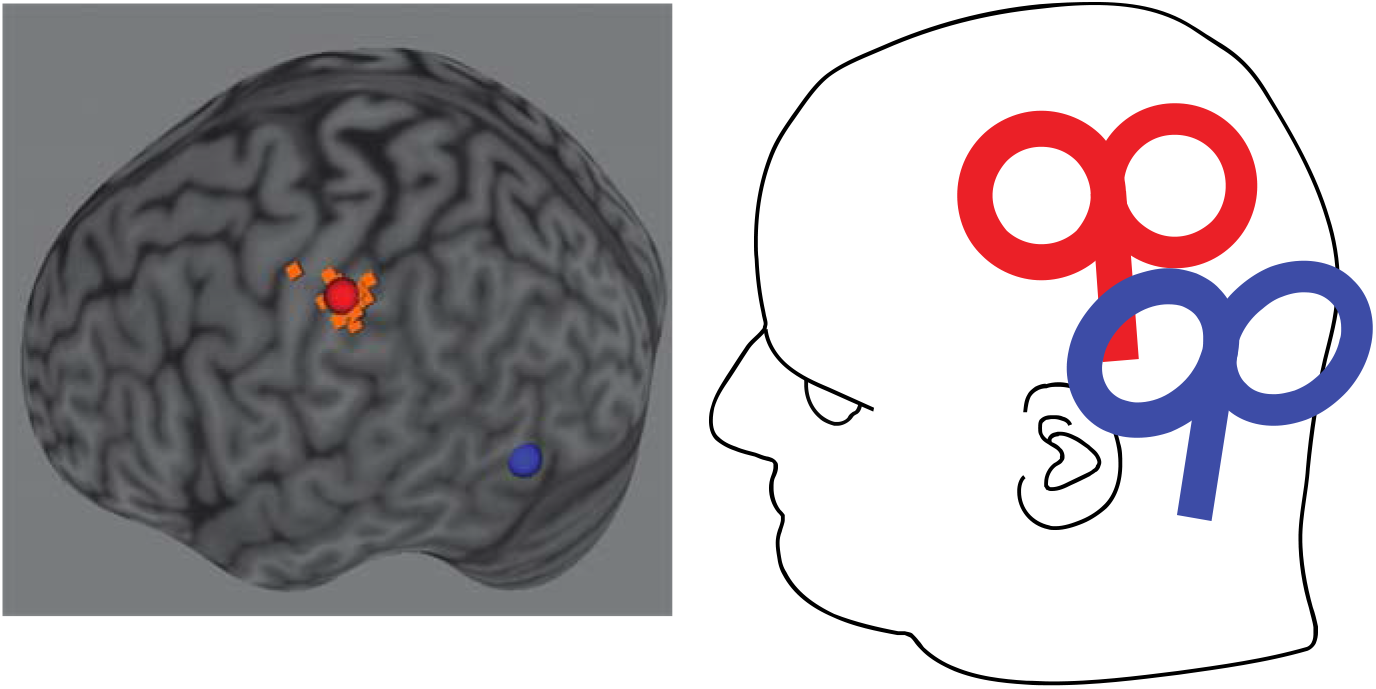
**A.** Brain areas targeted by TMS. Spheres show the mean location on a standard MNI brain (red: aIPS, blue: LO). Orange squares represent individual stimulation locations for aIPS. Note that for LO, coordinates from literature were used which were the same for all participants. **B**. Example of TMS coil positioning for aIPS (red) and LO (blue).

TMS was delivered with a 70mm figure-of-eight TMS DuoMag XT coil (Deymed Diagnostic). Electromyography (EMG) signals of the first dorsal interosseus (FDI) muscle were recorded with adhesive electrodes in a belly-tendon montage. The motor evoked potentials (MEPs) in response to TMS on the primary motor cortex (M1) were measured with the Brainsight system. In Brainsight, we defined the hotspot as the location on M1 that elicited the largest MEPs in the FDI muscle. The active motor threshold (aMT) was defined as the stimulation intensity when MEPs could be distinguished compared to background EMG in at least 5/10 stimulations while participants contracted the FDI muscle at submaximal levels. In the experiment, a stimulation level of 120% aMT was used. Average stimulation intensities were 48±7% and 53±9% of maximum stimulator output in the aIPS and LO group, respectively. We used the aMT and not the rest motor threshold (rMT) because participants would be stimulated in the experiment during active movements and not rest. MEPs are higher during muscle contraction (Buccolieri et al. 2004), resulting in aMTs usually being lower than rMTs (Buccolieri et al. 2004; Ngomo et al. 2012). Hence, using the aMT value would be more tuned to the active nature of the task and minimize the risk of spread towards other areas (e.g. M1 in the case of aIPS stimulation).

### 2.3 Task procedure

Two force sensors (Nano17, ATI Industrial Automation) were used to record lifting performance of the participants. Force data was sampled in 3 directions at a 1000 Hz frequency using a NI-USB 6343X (National Instruments, USA). The sensors were attached to a manipulandum (Figure 1a) which included a basket in which 3D printed cubes of different weights could be placed. The cubes were all of the same size (5 × 5 × 5 cm), but had different weights by filling them with different amounts of lead shot. A light (105 g) and a heavy (525 g) object were predominantly used in the experiment. To provide some variation in the weights, a dummy object of 317 g was presented in 10% of the trials. Finally, practice trials were performed with another object of 260 g. When the weight of the manipulandum (±120 g) was added, total weights of 2.2, 4.3, 6.3 and 3.8 N were obtained for the light, medium, heavy and practice object, respectively.

The participants were seated in front of a table. The manipulandum with the object was placed behind a switchable screen (Magic Glass) that could be in an opaque or a transparent state. In this way, the objects could be changed in between trials by the experimenter unseen by the participant. Participants were instructed to grasp and lift the object when the screen turned transparent and to hold it at a height of approximately 5 cm until the screen turned opaque again (±3 s) and then replace it back on the table (i.e. one ‘trial’). Participants were instructed to lift the object by placing the thumb and index finger on the force sensors. After they had replaced the object, they were asked to judge the weight of the object on a self-chosen scale with no constrained upper or lower limit.

During the participant’s movements, TMS was applied in 2/3^rd^ of the trials. Participants were instructed to ignore the TMS and continue their movement. TMS was always applied at 120% of aMT in a burst of 3 pulses at 10 Hz (total duration 200 ms). We used this burst of pulses to cover a larger time period of the lifting movement. Similar protocols with bursts of 2-5 pulses at 10 Hz were used in previous studies (Davare et al. 2006; Rice et al. 2006; White et al. 2013). Three TMS conditions were used: 1) TMS burst applied during dynamic loading, 2) during static holding or 3) no stimulation, which were presented in a pseudorandom order across the experiment. In the dynamic TMS condition (TMS_dynamic_) the first pulse was delivered when the participant contacted the object (i.e. grip force>0.4N). The three pulses together approximately covered the complete loading phase (see Figure 1B). In the static TMS condition (TMS_static_), TMS was applied during static object holding, where the first pulse was delivered 500 ms after lift-off (load force>object weight). In the no stimulation condition (TMS_no_), no TMS was applied during the trial. TMS triggering was controlled online based on sampled force data using a custom-written program in Labview (National Instruments) and Signal software (Cambridge Electronic Design Limited). A trigger was sent to the TMS stimulator from a personal computer through the NI-USB 6343X, which was in turn connected to a micro140-3 CED (Cambridge Electronic Design Limited).

Since we were interested in the effect of the previous lifted object on the present object, the order of object presentation was pseudo-randomized. Each object order (light-light, LL; heavy-light, HL; light-heavy, LH and heavy-heavy, HH) was presented 10 times for each stimulation condition in a random order. Therefore, 120 trials were used for analysis. In addition, 14 dummy trials (10% total) with a medium weight were presented. These trials, the first trial and the trials after dummy trials could not be analyzed because they had the wrong object order, which led to an extra 29 trials. For such trials, a stimulation condition was assigned randomly. In total, 149 trials were performed. Before the start of the experiment, the participant performed 5 practice trials to get familiar with the procedure, of which two trials included TMS (one in the dynamic phase, one in the static phase).

### 2.4 Data analysis

Trials where TMS was not applied correctly (i.e. before lifting, not at all) or the object was not lifted were removed from the perceptual analysis (n=16, 0.44%). In addition, trials where objects were lifted multiple times, dropped or when force data collection failed were also removed from the force analysis (n=25, 0.69%).

Participants’ weight estimates were converted to z-scores and averaged for each object order, TMS condition and TMS location.

Force data was filtered with a 2^nd^ order low-pass Butterworth filter with a cut-off frequency of 15 Hz. Grip force (GF) was defined as the mean of the horizontal forces of both sensors, while load force (LF) was the sum of the vertical forces. GF onset and LF onset were the time points at which the force crossed a threshold of 0.1 N. Note that the force threshold for triggering TMS_dynamic_ was slightly higher (0.4 N) to avoid responses to small initial bumps when grasping the object. Lift-off was the time point were LF overcame object weight. Grip force rate (GFR) and load force rate (LFR) were the differentiated forces. Since early force parameters are indicative of force planning, we were mostly interested in the peak force rates (Johansson and Westling 1988). The calculated force parameters are illustrated in Figure 1B. Peak GFR and peak LFR were defined as the highest peak values of the force rates between GF onset and 50 ms after lift-off. The time to peak force rate parameters (time to peak GFR and time to peak LFR) were calculated as the time between GF onset and the peak force rate.

While these parameters are typical of most force scaling studies, they all occur at early phases in lifting (i.e. before lift-off), so they cannot be affected by TMS_static_ which occurs after this time point. Therefore, we also calculated a late force parameter, which was static grip force (GF_static_). While during holding LF will normally not be much affected if the object does not move, the amount of GF can be more flexible while still maintaining a stable hold. We calculated GF_static_ as the average GF between 600 and 800 ms after lift-off, which was the period starting 100 ms after TMS_static_ was initiated (i.e. at the second pulse). This time period was chosen because it would be of the same duration as the stimulation period but was measured after the stimulation started so we could detect effects of stimulation on GF.

### 2.5 Statistical analysis

The variables of interest (perceptual answers, peak GFR, peak LFR, time to peak GFR, time to peak LFR) were analyzed with a mixed 2 (TMS location) × 3 (TMS condition) × 2 (previous weight) × 2 (current weight) analysis of variance (ANOVA). The factor TMS location was a between factor with the levels aIPS and LO for each participant group. The other factors were within factors, where TMS condition had three levels (TMS_dynamic_, TMS_static_, TMS_no_), and previous weight and current weight both had two levels (light or heavy). If a significant main effect or interaction with TMS location was found, the ANOVA was split into two 3 × 2 × 2 repeated measures ANOVAs to further analyze effects in both TMS groups. If Mauchly’s test indicated a violation of the sphericity assumption, a Greenhouse-Geisser correction was applied. A Bonferroni correction was used for post-hoc tests, which were performed with paired samples t-tests or independent t-tests for comparing within or between factors, respectively. The alpha-level was set at 0.05.

### 2.6 Relation between force scaling and perceptual estimates

To investigate the relation between motor planning and weight perception, we performed Pearson correlations between force rates (peak LFR and peak GFR), which are indicative of force planning (Johansson and Westling 1988), and perceptual estimates. We performed three types of correlations, to investigate effects of previous objects, TMS condition effects and trial - by-trial variations, respectively.

First, to test whether parameters were similarly affected by previous objects, we correlated sensorimotor memory effects with perceptual biases. We converted all variables to z-scores and subsequently subtracted values with a previous light object from values with a previous heavy object. This was done for each TMS location, TMS condition and for a current light and heavy object separately. Next, the obtained differences for perceptual estimates were correlated with the differences for peak LFR and peak GFR.

Second, we examined whether force and perceptual parameters were similarly affected by TMS over a specific location. For this, we used the z-scored variables and subtracted the baseline condition (TMS_no_) from the other TMS conditions (TMS_dynamic_ and TMS_static_) for each TMS location and current object weight separately. Because we did not find TMS condition interactions with previous weight (see Results), we pooled over previous object weight before calculating differences. We correlated the differences for the perceptual estimates with those for peak LFR and peak GFR.

Finally, we performed trial-by-trial correlations. For each participant, we correlated the trials of the perceptual estimates with peak LFR and peak GFR, for each TMS location, TMS condition and object weight separately. To see whether R-values were significantly different from zero, we performed one-sample t-tests for each condition.

## 3 RESULTS

We investigated the role of aIPS and LO in force scaling and weight perception when lifting objects. TMS was applied in the dynamic loading phase (TMS_dynamic_), the static holding phase (TMS_static_) or not at all (TMS_no_, control condition). The results for the perceptual estimates and force parameters are shown in Figure 3 and Table 1. Average force traces for each object, and each TMS location and TMS condition are shown in Figure 4. Differences with respect to the no stimulation condition (TMS_no_) are illustrated in Figure 5, to more clearly indicate the effects of stimulation.

**Figure 3.**
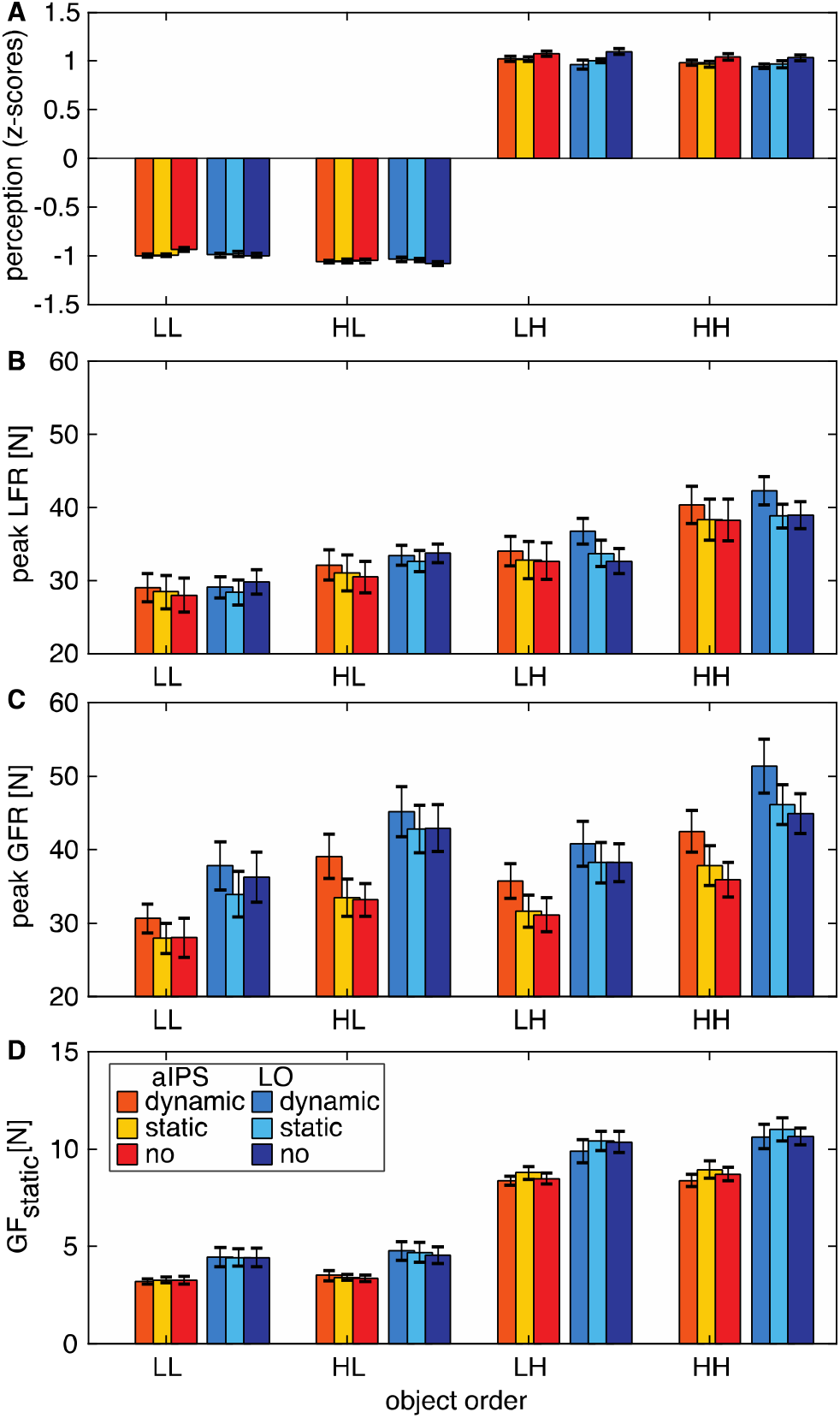
Results for **A** perceptual estimates, **B** peak load force rates (peak LFR), **C** peak grip force rates (peak GFR), and **D** grip force during static holding (GF_static_) for each object order (light-light: LL, heavy-light: HL, light-heavy: LH, heavy-heavy: HH). Colors indicate TMS group (red shades: aIPS, blue shades: LO) for the stimulation conditions: TMS_dynamic_, TMS_static_, TMS_no_ (control condition). Error bars represent standard errors. Note that a main effect of previous weight was found for all parameters, that did not interact with TMS location or condition (mixed ANOVA). N=15.

**Table 1.** Results for all parameters: perception, peak load force rate (peak LFR), peak grip force rate (peak GFR), grip force during static holding (GF_static_), time to peak LFR (tpLFR) and time to peak GFR (tpGFR). The upper and lower table shows values for the aIPS and LO group, respectively. Columns are separated for object order (light-light: LL, heavy-light: HL, light-heavy: LH, heavy-heavy: HH), and TMS condition TMS_dynamic_, TMS_static_, TMS_no_). Values represent means ± SEM.

| aIPS | LL |  |  | HL |  |  | LH |  |  | HH |  |  |
| --- | --- | --- | --- | --- | --- | --- | --- | --- | --- | --- | --- | --- |
|  | dynamic | static | no | dynamic | static | no | dynamic | static | no | dynamic | static | no |
| perception | -1.00±0.01 | -1.00±0.01 | -0.93±0.02 | -1.06±0.01 | -1.05±0.02 | -1.05±0.02 | 1.02±0.02 | 1.02±0.02 | 1.07±0.03 | 0.98±0.03 | 0.97±0.03 | 1.04±0.03 |
| Peak LFR | 29.02±1.96 | 28.44±2.30 | 28.01±2.34 | 32.15±2.07 | 31.07±2.43 | 30.47±2.15 | 34.06±2.01 | 32.79±2.58 | 32.64±2.49 | 40.37±2.57 | 38.36±2.79 | 38.28±2.84 |
| Peak GFR | 30.64±1.95 | 27.95±2.06 | 28.04±2.66 | 39.11±2.97 | 33.49±2.50 | 33.22±2.22 | 35.79±2.38 | 31.67±2.18 | 31.17±2.33 | 42.48±2.84 | 37.84±2.66 | 35.92±2.36 |
| $GF_{static}$ | 3.20±0.13 | 3.29±0.15 | 3.27±0.20 | 3.52±0.27 | 3.41±0.15 | 3.38±0.16 | 8.38±0.24 | 8.79±0.33 | 8.49±0.29 | 8.40±0.32 | 8.95±0.44 | 8.72±0.35 |
| tpLFR | 0.17±0.01 | 0.16±0.01 | 0.16±0.01 | 0.17±0.01 | 0.17±0.01 | 0.17±0.01 | 0.24±0.02 | 0.23±0.02 | 0.23±0.02 | 0.23±0.01 | 0.23±0.02 | 0.22±0.02 |
| tpGFR | 0.18±0.01 | 0.17±0.01 | 0.18±0.02 | 0.19±0.01 | 0.18±0.02 | 0.19±0.01 | 0.25±0.02 | 0.26±0.03 | 0.25±0.03 | 0.23±0.02 | 0.24±0.02 | 0.23±0.02 |

| LO | LL |  |  | HL |  |  | LH |  |  | HH |  |  |
| --- | --- | --- | --- | --- | --- | --- | --- | --- | --- | --- | --- | --- |
|  | dynamic | static | no | dynamic | static | no | dynamic | static | no | dynamic | static | no |
| perception | -0.99±0.02 | -0.98±0.03 | -1.00±0.02 | -1.04±0.02 | -1.04±0.02 | -1.08±0.02 | 0.96±0.05 | 1.00±0.02 | 1.10±0.03 | 0.95±0.03 | 0.97±0.04 | 1.03±0.03 |
| Peak LFR | 29.07±1.46 | 28.40±1.70 | 29.84±1.69 | 33.46±1.36 | 32.66±1.45 | 33.77±1.28 | 36.77±1.80 | 33.71±1.78 | 32.67±1.74 | 42.28±1.93 | 38.86±1.63 | 38.94±1.86 |
| Peak GFR | 37.80±3.25 | 33.95±3.06 | 36.28±3.40 | 45.18±3.39 | 42.80±3.25 | 42.94±3.17 | 40.81±3.07 | 38.27±2.77 | 38.28±2.58 | 51.36±3.65 | 46.15±2.70 | 44.94±2.70 |
| $GF_{static}$ | 4.46±0.49 | 4.43±0.45 | 4.44±0.47 | 4.78±0.48 | 4.70±0.50 | 4.55±0.43 | 9.90±0.59 | 10.42±0.49 | 10.37±0.54 | 10.63±0.62 | 11.02±0.60 | 10.66±0.44 |
| tpLFR | 0.12±0.01 | 0.12±0.01 | 0.12±0.01 | 0.12±0.01 | 0.12±0.01 | 0.12±0.01 | 0.20±0.01 | 0.18±0.02 | 0.17±0.01 | 0.19±0.01 | 0.16±0.01 | 0.17±0.01 |
| tpGFR | 0.15±0.01 | 0.13±0.01 | 0.14±0.01 | 0.15±0.01 | 0.15±0.01 | 0.15±0.01 | 0.23±0.02 | 0.22±0.02 | 0.20±0.02 | 0.21±0.01 | 0.19±0.01 | 0.19±0.02 |

**Figure 4.**
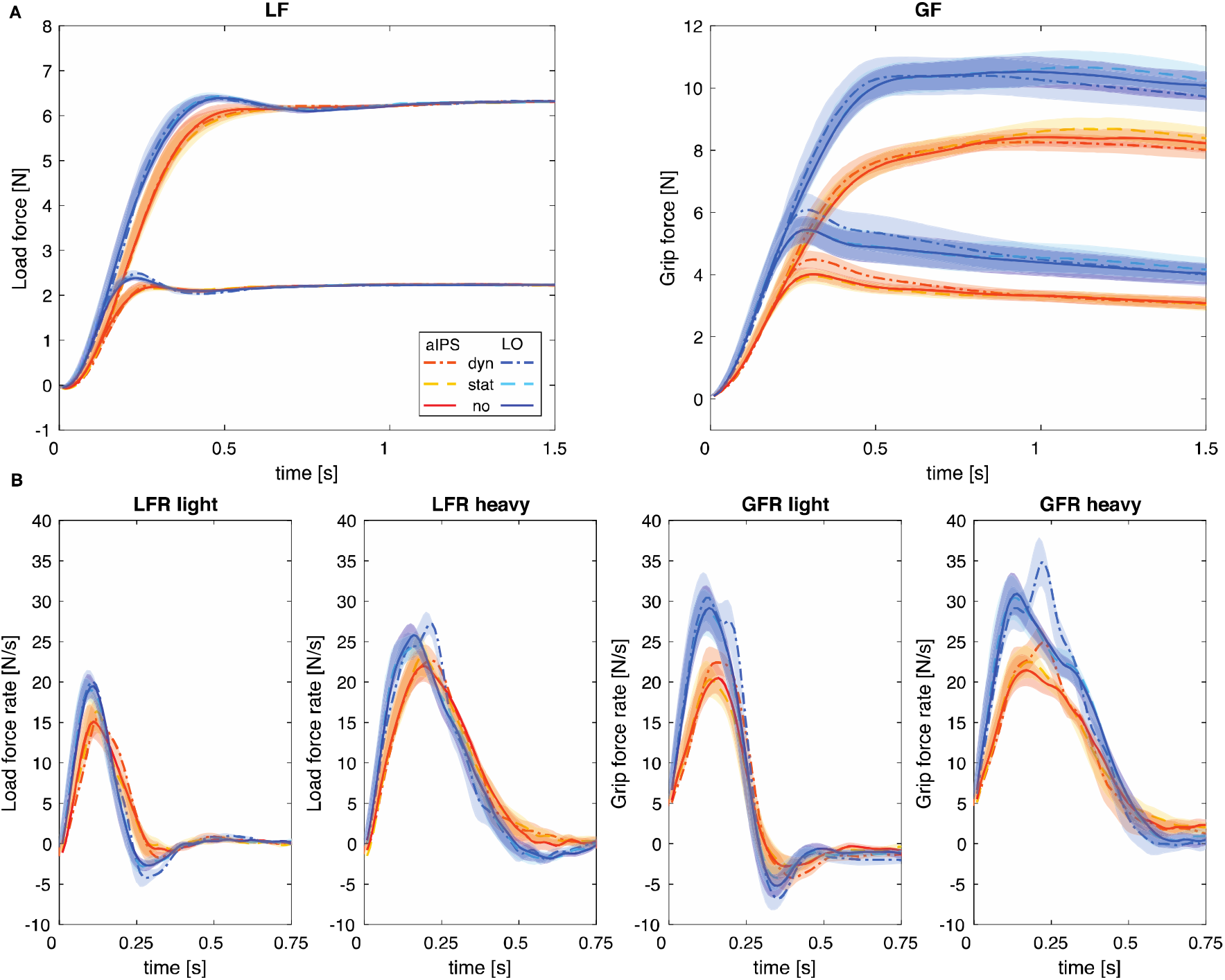
**A.** Average force traces of load forces (LF) and grip forces (GF). The upper traces are for heavy objects, the lower traces for light objects. **B.** Average traces of force rates (LFR and GFR) for light and heavy objects separately. Lines represent TMS conditions (dash-dot, ‘dyn’: TMS_dynamic_; dashed, ‘stat’: TMS_static_; solid, ‘no’: TMS_no_) for the aIPS group (red shades) and the LO group (blue shades). Shadings indicate standard errors. N=15.

**Figure 5.**
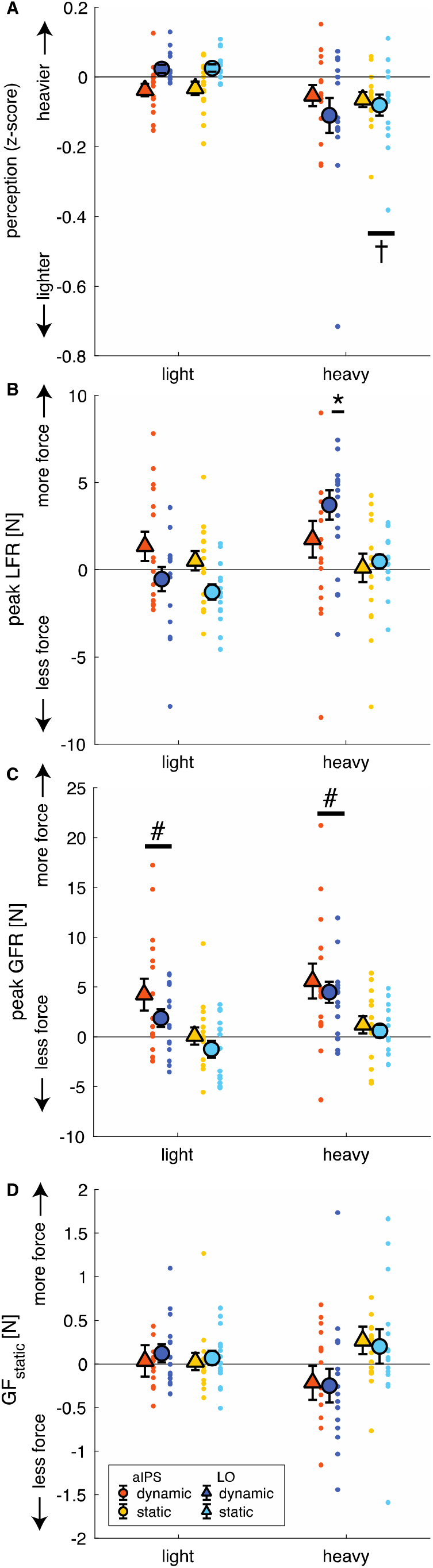
Stimulation effects displayed as differences with respect to the TMS_no_ condition. Panels indicate results for **A** perceptual estimates, **B** peak load force rates (peak LFR), **C** peak grip force rates (peak GFR) and **D** grip force during static holding (GF_static_) for light and heavy objects separately. Note that there is pooled over object order, since TMS did not interact with the effect of previous weight. Colors and symbols indicate TMS group (triangles: aIPS; circles: LO) for the stimulation conditions (TMS_dynamic_ and TMS_static_). Error bars represent standard errors, dots indicate individual subjects. †significant effect for TMS_static_, regardless of TMS location. *significant effect for TMS_dynamic_ for LO only. #significant effect for TMS_dynamic_, regardless of TMS location and object weight. N=15.

### 3.1 TMS affects weight perception but not the perceptual bias

#### Behavioral effects

The ANOVA on the perceptual weight estimates showed the expected main effect of current weight (F(1,28)=66786.5, p<0.001, *η*_p_^2^=1.00) where light objects were rated lighter than heavy objects (Figure 3). In addition, the effect of previous weight (F(1,28)=11.7, p=0.002, *η*_p_^2^=0.29) indicated that a perceptual bias was observed, where an object was perceived to be heavier when the previously lifted object was light (0.023±0.01) compared to when it was heavy (−0.032±0.01). This effect corroborates our previous findings, indicating a perceptual bias from previous lifted objects (van Polanen and Davare 2015b).

#### TMS effects

The effect of previous weight did not interact with TMS condition or with TMS location, suggesting that the weight perception bias from the preceding object was not affected by TMS. However, we did observe a main effect of TMS condition (F(1.5,41.9)=6.3, p=0.008, *η*_p_^2^=0.18) and an interaction of current weight × TMS condition (F(1.7,46.5)=6.1, p=0.007, *η*_p_^2^=0.18). Effects of stimulation are shown in Figure 5A and show that weight estimation was lower after stimulation during both stimulation conditions compared to TMS_no_. However, these results should be interpreted in light of the interaction of current weight × TMS condition. Post-hoc tests of the interaction indicated that perceptual ratings were only lower after TMS_static_ compared to TMS_no_ and only for heavy objects (p=0.004).

Since there was also a trend for a TMS location × TMS condition × current weight interaction (F(1.67,46.5)=3.1, p=0.065, *η*_p_^2^=0.10), we further explored this interaction by performing two separate repeated measures ANOVAs on the TMS locations to shed light on these stimulation effects. For aIPS, we only found main effects (current weight: F(1,14)=58349.6, p<0.001, *η*_p_^2^=1.00; previous weight: F(1,14)=6.9, p=0.020, *η*_p_^22^=0.33; TMS condition: F(2,28)=5.0, p=0.014, *η*_p_^2^=0.26). Similar to the main effects of the mixed ANOVA, these effects indicated that objects were perceived as lighter when current objects were (a) light compared to heavy, (b) lifted after heavy objects compared to light objects, and (c) after TMS_static_ compared to TMS_no_. No interaction effects were found in the aIPS group. However, for the LO group, we found that the current weight × TMS condition effect was significant as well, in addition to the main effects of current weight and previous weight (current weight: F(1,14)=23310.3, p<0.001, *η*_p_^2^=1.00; previous weight: F(1,14)=4.8, p=0.046, *η*_p_^2^=0.26; current weight × TMS condition: F(2,28)=6.1, p=0.007, *η*_p_^2^=0.30). Further post-hoc tests for this interaction revealed no significant effects of TMS condition after Bonferroni corrections. No significant differences were found between the two TMS groups on any condition or current weight condition. Overall, although the perceptual bias induced by object order was not affected by TMS, stimulation in the static phase, over either aIPS or LO, seemed to affect weight perception.

### 3.2 Early force parameters: LO stimulation affects load force scaling

To test effects of TMS on parameters in the early phases of lifting (dynamic phase), we looked at the peak values of the force rates. Results for peak LFR are shown in Figure 3B. The mixed ANOVA on peak LFR revealed main effects of current weight (F(1,28)=171.8, p<0.001, *η*_p_^2^=0.86), previous weight (F(1,28)=208.5, p<0.001, *η*_p_^2^=0.88) and TMS condition (F(1.3,35.2)=9.8, p=0.002, *η*_p_^2^=0.26). In addition, interactions of current weight × previous weight (F(1,28)=9.0, p=0.006, *η*_p_^2^=0.24), current weight × TMS condition (F(2,56)=4.9, p=0.010, *η*_p_^2^=0.15) and a triple interaction of current weight × TMS condition × TMS location (F(2,56)=3.2, p=0.048, *η*_p_^2^=0.10) were found.

#### Behavioral effects peak LFR

To start with the current weight × previous weight interaction, post-hoc tests showed that all comparisons were significant (Figure 3B). Peak LFR was lower for light objects compared to heavy objects (both previous weights: p<0.001) and previous light objects were lifted with a lower peak LFR compared to previous heavy weights (both objects: p<0.001). These results indicate that force scaling was based on the current weight and on previously lifted objects, both for light and heavy weights. Perhaps the interaction effect could be explained by the notion that the effect of previous weight was somewhat stronger in heavy objects than light objects.

#### Effects of aIPS TMS

To further investigate the triple interaction, we performed separate repeated measures ANOVAs for the two TMS locations. No significant differences were found between the two TMS groups for any of the object weights or TMS conditions. In the separate ANOVA for the aIPS group, main effects of current weight (F(1,14)=63.2, p<0.001, *η*_p_^2^=0.82), previous weight (F(1,14)=83.9, p<0.001, *η*_p_^2^=0.86) and an interaction of current weight × previous weight (F(1,14)=6.2, p=0.026, *η*_p_^2^=0.31) were found. These results were the same as those found in the mixed ANOVA (Figure 3B), indicating that peak LFR was affected both by the current and by the previous object. Stimulation effects, i.e. differences with respect to the TMS_no_ condition, are shown in Figure 5B. The absence of effects of TMS condition suggests that TMS over aIPS did not influence peak LFR.

#### Effects of LO TMS

For the LO group, we also found main effects of current weight (F(1,14)=140.0, p<0.001, *η*_p_^2^=0.91) and previous weight (F(1,14)=129.1, p<0.001, *η*_p_^2^=0.90). However, for the LO group, we also found effects of TMS condition (F(1.2,16.7)=10.3, p=0.004, *η*_p_^2^=0.42) and a current weight × TMS condition interaction (F(2,28)=9.6, p=0.001, *η*_p_^2^=0.41). Post-hoc tests for this double interaction showed that in the TMS_dynamic_ condition, a higher peak LFR was seen than in the TMS_static_ or TMS_no_ condition, but only for heavy objects (both conditions, p<0.014, see Figure 5B). In other words, heavy objects were lifted with higher load force rates when TMS was applied in the dynamic phase. The peak LFR differed between light and heavy objects for all TMS conditions (all p<0.001).

#### Effects on time to peak LFR

For time to peak LFR, main effects of current weight (F(1,28)=246.7, p<0.001, *η*_p_^2^=0.90), TMS condition (F(2,56)=3.5, p=0.037, *η*_p_^2^=0.11) and TMS location (F(1,28)=8.7, p=0.006, *η*_p_^2^=0.24) were found. In addition, there were interactions of current weight × TMS condition (F(1.6,46.7)=4.9, p=0.016, *η*_p_^2^=0.15) and current weight × previous weight (F(1,28)=4.9, p=0.035, *η*_p_^2^=0.15). The main effect of location showed a longer time to peak LFR in the aIPS group compared to the LO group. Because of this main effect of location, we split the ANOVAs for the two TMS groups. For aIPS, only a main effect of current weight was found (F(1,14)=115.0, p<0.001, *η*_p_^2^=0.89), indicating that the time to peak LFR was longer for heavy objects compared to light ones (Table 1). For LO, main effects of current weight (F(1,14)=136.6, p<0.001, *η*_p_^2^=0.91), TMS condition (F(2,28)=4.1, p=0.028, *η*_p_^2^=0.23) and an interaction of current weight × TMS condition (F(2,28)=6.9, p=0.004, *η*_p_^2^=0.33) were found. Similar to the effect of current weight in the aIPS group, post-hoc tests indicated that peak LFR occurred later when lifting heavy objects than light objects in all stimulation conditions (all p<0.001). Interestingly, there was also an effect of TMS condition, where the time to peak LFR was longer after TMS_dynamic_ compared to TMS_no_ when lifting heavy objects (p=0.009). This later LFR peak after stimulation in the dynamic phase is also visible in Figure 4B (second panel).

To summarize, it appears that TMS over aIPS did not influence load force scaling. On the other hand, LO stimulation delivered during the dynamic phase increased peak force rates, and their time to peak, only for heavy objects. When one looks at the force profiles in Figure 4, it can be seen that after LO stimulation, an extra peak is visible in the LF rates (Figure 4B, LFR heavy), which could account for the increased maximum value at a later time point.

### 3.3 Early force parameters: Grip force rates are affected by TMS, but not specifically for TMS location

#### Behavioral effects on peak GFR

As expected, peak GFR was higher after lifting a previous heavy weight compared to a previous light weight (effect previous weight: F(1,28)=204.9, p<0.001, *η*_p_^2^=0.88; Figure 3C). In addition, effects of current weight (F(1,28)=44.6, p<0.001, *η*_p_^2^=0.61), TMS condition (F(1.5,41.1)=18.4, p<0.001, *η*_p_^2^=0.40) and an interaction of current weight × TMS condition (F(2,56)=3.6, p=0.035, *η*_p_^2^=0.11) were found. Heavy objects were lifted with higher grip force rates than light objects in all TMS conditions (all p<0.024). In accordance with the results on peak LFR, this indicates that grip forces were scaled according to both current and previous object weights.

#### TMS effects on peak GFR

Furthermore, both for light and heavy objects, a higher peak GFR was seen after TMS_dynamic_ compared to TMS_static_ and TMS_no_ (all p<0.023, Figure 5C). The interaction effect of current weight × TMS condition could be explained by somewhat larger stimulation effects for heavy than light objects. A main effect of location was also found (F(1,28)=4.4, p=0.046, *η*_p_^2^=0.14), but separate ANOVAs showed the same main effects (all F>8.0, all p<0.003) for both TMS groups with no significant interaction effects. This indicated that there was just an overall difference in grip force rate that was higher in the LO group (Figure 3C), independently of TMS condition. Therefore, the peak GFR was increased when TMS was applied in the dynamic phase for both TMS groups.

#### Effects on time to peak GFR

The results for the time to peak GFR are shown in Table 1. The mixed ANOVA on the time to peak GFR showed a main effect of current weight (F(1,28)=60.0, p<0.001, *η*_p_^2^=0.68) and an interaction of current weight × previous weight (F(1,28)=16.4, p<0.001, *η*_p_^2^=0.37). Post-hoc tests indicated that heavy objects had a longer time to peak GFR than light objects (both previous weights: p<0.001). However, for light objects, a previous light weight resulted in an earlier peak GFR compared to a previous heavy lift (p=0.007), whereas this was reversed for heavy objects (p=0.013). No effects nor interactions with TMS condition or TMS location were found, indicating that the time to peak GFR was not affected by TMS.

All in all, it seems that TMS in the dynamic phase affected the magnitude of the peak grip force rate, but not the latency between force onset and peak GFR. Also, the effect of previous weight on GFR was not altered by TMS. Since both LO and aIPS stimulation had the same effect on grip force rates, this suggests a non-specific TMS effect rather than an actual indication of LO or aIPS contribution to grip force scaling. Because of these results, we also looked at TMS effects on grip force in the later phases of lifting to see whether these would also be similarly affected by TMS on both locations.

### 3.4 Late force parameter: no effects of TMS on static grip force

Since the other force parameters all occurred before TMS_static_, they could not be affected by this stimulation during the static phase. Therefore, we also investigated GF_static_, which was the grip force during static holding. Main effects of current weight (F(1,28)=1276.0, p<0.001, *η*_p_^2^=0.98), previous weight (F(1,28)=12.7, p=0.001, *η*_p_^2^=0.31) and TMS location (F(1,28)=8.8, p=0.006, *η*_p_^2^=0.24) were found. In addition, interaction effects of current weight × TMS condition (F(2,56)=7.5, p=0.001, *η*_p_^2^=0.21), current weight × TMS location (F(1,0)=4.5, p=0.044, *η*_p_^2^=0.14) and a triple interaction of current weight × previous weight × TMS location (F(1,0)=5.3, p=0.029, *η*_p_^2^=0.16) were found.

#### Effect of aIPS TMS

When the ANOVA was split for both TMS groups, only a main effect of current weight (F(1,14)=757.4, p<0.001, *η*_p_^2^=0.98) and an interaction of current weight × TMS condition (F(1.4,19.5)=4.6, p=0.034, *η*_p_^2^=0.25) were found for the aIPS group. This indicated that GF_static_ was higher for heavy objects than light ones (all conditions: p<0.001, Figure 3D), but post-hoc tests did not show significant TMS condition effects. When observing Figure 4A (right panel), the interaction might be explained by lower GF values after dynamic stimulation and higher values after static stimulation, for heavy objects only. However, these stimulation results were not statistically significant (Figure 5D).

#### Effects of LO TMS

For the LO group, main effects of current weight (F(1,14)=570.6, p<0.001, *η*_p_^2^=0.98), previous weight (F(1,14)=31.8, p<0.001, *η*_p_^2^=0.70) and an interaction of current weight × previous weight (F(1,14)=15.7, p=0.001, *η*_p_^2^=0.53) were found. Post-hoc tests revealed that GF_static_ was higher for heavy objects compared to light ones (both previous weights p<0.001) and higher when a heavy object was previously lifted compared to a previous light lift (both current weights p<0.008). There were no effects of TMS condition, indicating that LO stimulation did not influence GF_static_. Finally, further tests for the effect of TMS location revealed that the TMS groups differed significantly in the LH (p=0.036) and HH (p=0.012) conditions, indicating that higher grip forces were used by the LO group. Since TMS location did not interact with TMS condition, this suggest a general group difference.

Since the effect of previous weight was only found in the LO group and not the aIPS group, this could suggest that aIPS stimulation eradicated the order effect on grip forces. However, since there was no interaction with TMS condition, indicating that this effect was also absent in TMS_no_ in the aIPS group, this result might merely reflect a general group difference without any stimulation effects.

### 3.5 Correlations between force scaling and perceptual estimates

Because we found both effects of previous object weight for force parameters and perceptual estimates (see above), we examined whether these effects were related. We correlated effects of previous object weight for the perceptual estimates with peak LFR and peak GFR, which are shown in Table 2. Only the correlation of perception with peak LFR in the aIPS group for the TMS_dynamic_ condition when lifting a light object was significant (R=-0.60, p=0.017; see Figure 6A). This correlation indicates that larger sensorimotor memory effects are associated with larger perceptual biases. Since no other correlations were significant, and none in the TMS_no_ condition, the association between order effects on force scaling and perceptual estimates appears weak.

**Table 2.** Between-subject correlations (R-values) of sensorimotor memory and perceptual bias effects. Differences between previous heavy and previous light were calculated for z-score force parameters and correlated with the same differences of z-scored perceptual estimates, for light and heavy lifts and each TMS condition (dynamic, static, no) separately. Force parameters are peak load force rate (peak LFR), peak grip force rate (peak GFR). *p<0.05.

|  | aIPS |  |  |  |  |  | LO |  |  |  |  |  |
| --- | --- | --- | --- | --- | --- | --- | --- | --- | --- | --- | --- | --- |
|  | light |  |  | heavy |  |  | light |  |  | heavy |  |  |
|  | dynamic | static | no | dynamic | static | no | dynamic | static | no | dynamic | static | no |
| Peak LFR | -0.60* | 0.15 | 0.20 | -0.03 | -0.28 | -0.25 | -0.23 | -0.31 | -0.24 | -0.27 | 0.09 | -0.04 |
| Peak GFR | -0.43 | 0.10 | 0.18 | 0.23 | -0.40 | -0.18 | -0.10 | -0.09 | -0.14 | -0.11 | -0.09 | -0.10 |

**Figure 6.**
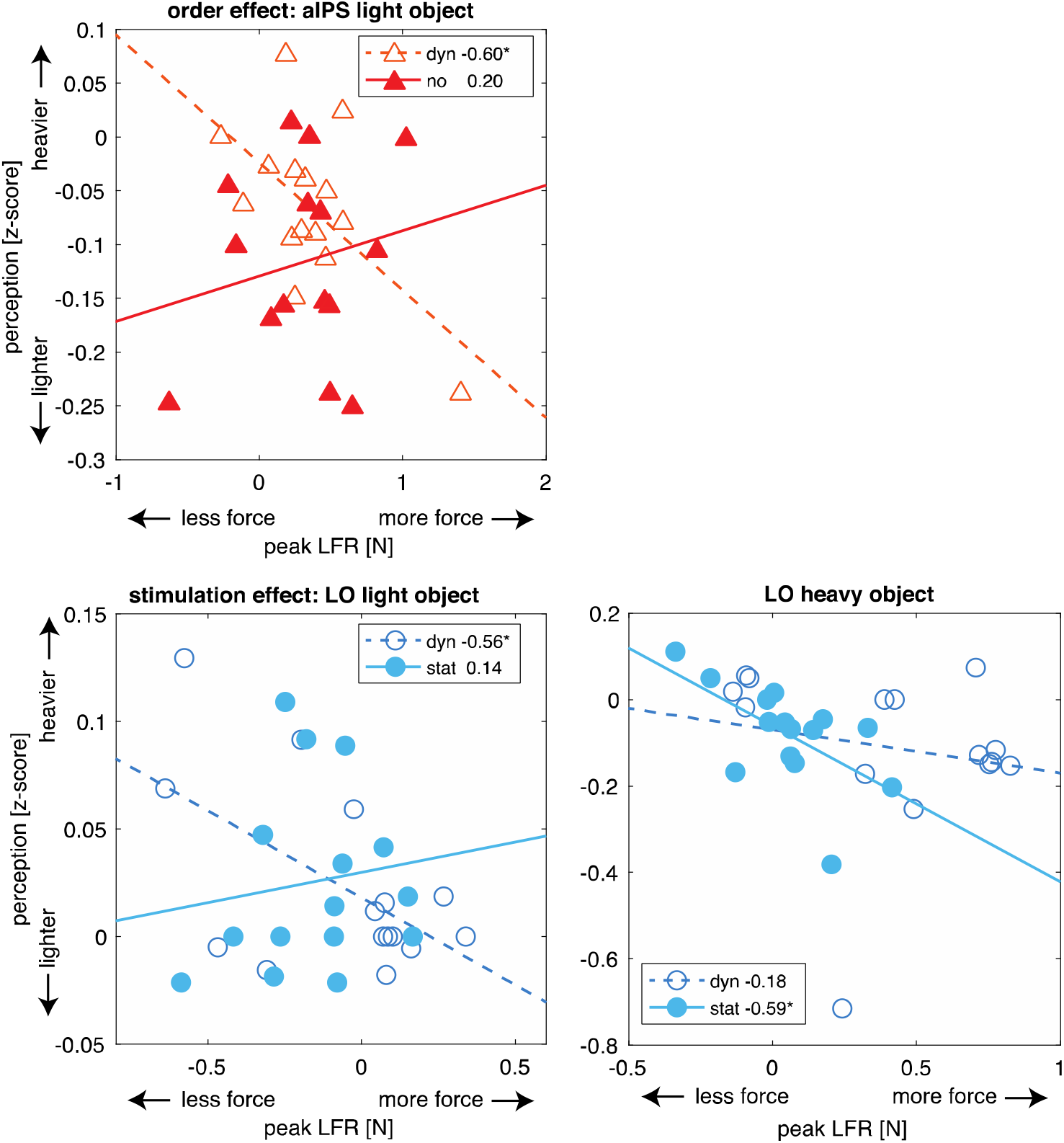
**A.** Correlations between order effects (previous heavy-previous light) for peak load force rates (peak LFR, sensorimotor memory effects) and perception (perceptual bias) for the aIPS group, for the light object. A significant correlation was found for the TMS_dynamic_ (dyn), but not for the TMS_no_ condition (no). B. Correlations between stimulation effects (stimulation-TMS_no_) for peak LFR and perception. Significant correlations were found for TMS_dynamic_ (dyn), but not TMS_static_ (stat) for a light object (left panel). Significant correlations were found for TMS_static_, but not TMS_dynamic_ for a heavy object (right panel). Symbols indicate individual participants (N=15). Legends also display R-values. Y-axes indicate whether objects are perceived to be heavier or lighter after lifting a heavy object. X-axes indicate whether objects were lifted with more or less force after lifting a heavy object. *p<0.05.

To test whether effects of TMS on force parameters were related to TMS effects on perceptual estimates, we correlated stimulation effects for these variables. Since we did not find stimulation effects that interacted with previous object effects, we averaged over object order to obtain means for the light and heavy object separately. Differences with respect to the TMS_no_ condition were correlated and these are shown in Table 3. Only significant correlations were seen in the LO group (see Figure 6B), where peak LFR correlated with perception in the TMS_dynamic_ condition for light objects (R=-0.56, p=0.029) and in the TMS_static_ condition for heavy objects (R=-0.59, p-0.019). No correlations were significant for peak GFR. These negative correlations for the LO group suggest that increases in peak LFR due to TMS are associated with decreases in weight perception. However, since stimulation in the static phase cannot affect peak LFR because it occurs after the measurement of this parameter, this correlation is unexpected. It is possible that the parameters, but not the stimulation effects, are related. In other words, variations in peak LFR could be related to variations in perceptual estimates. Therefore, we also correlated trial-by-trial variations for each participant and tested whether R-values were different from zero. Here we only found a significant effect for correlations between peak LFR and perception, not between peak GFR and perception. More specifically, only the R-values for the LO group in the TMS_dynamic_ condition when lifting light objects were significantly different from zero (R=-0.13±0.05, t(14)=-2.53, p=0.024). Overall, correlations were mainly found for the LO group, between peak LFR and weight perception. However, the correlations between forces and estimates were primarily observed for the light objects, whereas the results showed stimulation effects on weight perception and peak LFR for heavy objects only. Therefore, it is not clear whether stimulation had similar effects on both parameters or whether the correlations reflect a general relation between the parameters.

**Table 3.**
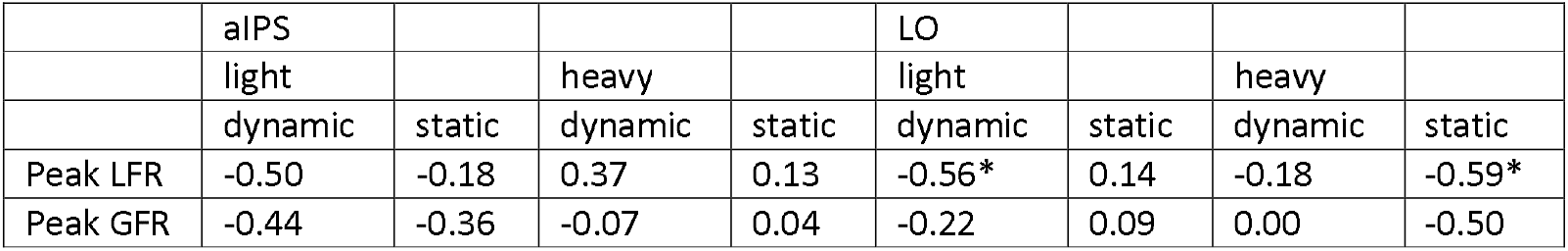
Between-subject correlations (R-values) of stimulation effects (compared to no-stimulation) between force parameters and perceptual estimates. Z-scored variables were averaged over object order, to obtain values for light and heavy objects, and differences with respect to the no stimulation (TMS_no_) were calculated, for TMS_dynamic_ and TMS_static_ separately. Force parameters are peak load force rate (peak LFR), peak grip force rate (peak GFR). *p<0.05.

## 4 DISCUSSION

In a recent study, we showed that sensorimotor memory effects were related to weight estimations (van Polanen and Davare 2015b). That is, both force scaling and weight perception were affected by the previously lifted object weight. Moreover, these effects correlated both across participants and in within-participant trial-by-trial comparisons. In the present experiment, we wanted to investigate the neural underpinnings of these effects in object lifting and weight perception. We hypothesized a role for aIPS, known to be involved in force scaling (Dafotakis et al. 2008; Davare et al. 2007) and LO, which is important in object perception (Amedi et al. 2001), and used TMS to infer their causal role. Whereas we replicated the effects of object order both for force scaling and weight perception, we did not find strong associations between action and perception components. Furthermore, although we did find effects of both aIPS and LO stimulation on force scaling and weight perception, these stimulation effects did not alter the effect of previously lifted objects on force scaling and weight perception. Therefore, although these areas might play a role in object lifting and weight perception, they do not seem to mediate effects of force planning based on previous experience on lifting performance and weight estimation. In the next paragraphs, we will further elaborate on these findings.

First of all, we did replicate effects of object order on force scaling and weight perception. When a heavy object was previously lifted, force scaling was larger than when the previously lifted object was light. This corroborates studies showing that forces are planned based on the sensorimotor memory of previous objects (Gordon et al. 1994; Johansson and Westling 1988; Loh et al. 2010; van Polanen and Davare 2015b). In addition, we found that when previously lifting a heavy object, the present object felt lighter than when a light object was previously lifted, replicating the perceptual bias found in previous studies (Maiello et al. 2018; van Polanen and Davare 2015b). In our previous study, we only found an effect for current light objects, not heavy (van Polanen and Davare 2015b). We argued in that study that for heavy objects differences needed to be larger to be perceptually discriminable. The findings in this study indicate that the perceptual bias can be seen for both light and heavy objects since we found no interaction of previous weight with current weight. Since the present study has a larger power (N=30) compared to the previous study (N=10), this might explain why we could detect an effect for both object weights.

In contrast to our previous study, we only found few correlations between force scaling parameters and perception, both for comparisons across participants and within-participant trial-by-trial comparisons. Although few studies have compared these order effects on action and perception components (Schneider et al. 2019; van Polanen and Davare 2015b), the lack of correlation in this study casts doubt on the hypothesis that these effects stem from a common underlying mechanism. It would suggest that the relation is weak or not very robust. More specifically, the TMS procedure could have weakened the relation between force control and perceptual effects. Further research is needed to provide more insight into the involved processes. In the present study, we investigated the role of the aIPS and LO, but these areas do not seem to mediate effects of previous lifting experience on force lifting or weight perception, nor on the potential association of these effects.

Several other areas could be proposed to play a role in force planning based on previous experience. It has been well-established that sensorimotor memory is represented in the primary motor cortex (M1) (Berner et al. 2007; Chouinard et al. 2005; Loh et al. 2010; Nowak et al. 2005). However, it is likely that M1 receives input from other areas (Parikh et al. 2014). For instance, it has been shown that effects on grip force scaling can be differently affected depending on the timing of M1 stimulation (Berner et al. 2007). Furthermore, it is known that inputs to M1 change based on grasp type, through connections with the ventral premotor area (PMv) and indirectly from aIPS (Davare et al. 2007; Davare et al. 2008). For these areas, it was shown that PMv plays a role in predictive force scaling according to recent lifts, whereas aIPS is involved in force adjustments (Dafotakis et al. 2008). Functional MRI studies have shown more areas that seem to be involved in unpredictable weight changes in fronto-parietal circuits (Schmitz et al. 2005), but also the primary somatosensory cortex (Jenmalm et al. 2006). Therefore, it looks like a network of areas is involved in force planning according to previous weight experience. Possibly, aIPS is more concerned with online corrections of movements, as has been observed in grasping tasks (Glover et al. 2005; Rice et al. 2006; Tunik et al. 2005) or force adjustments in lifting tasks (Dafotakis et al. 2008), but less with predictive scaling according to previous experience.

The neural network of weight perception is less clear from the literature. Some studies indicate that weight is represented in LO (Gallivan et al. 2014). Other studies suggest roles for M1 in representing weight (Chouinard et al. 2009) and sense of effort in force production (Takarada et al. 2014). However, it is not exactly clear whether these findings also hold for judging weight. The present findings indicate that although aIPS and LO did not seem to be involved in the perceptual bias from previous lifted objects, an effect on weight perception by TMS, independent of stimulation site, was observed. When TMS stimulation was applied during the static phase, heavy objects were judged to be lighter. Since there are connections between parietal and temporal areas (Budisavljevic et al. 2018; Ramayya et al. 2010), it is possible that perceptual processing of weight runs through connections between these areas. Interestingly, a trend was observed for a specific object weight effect for LO stimulation: if LO was stimulated, heavy objects appeared to be lighter, whereas light objects were not perceived differently. Such a specific weight effect was not seen after aIPS stimulation. This result could be interpreted as a decrease in weight discrimination, where the different objects could be less well discriminated in weight, i.e. making heavy objects appear lighter and light objects appear heavier. This would indicate that LO is involved in weight discrimination. Although this is a speculative conclusion from our study results, this is an interesting notion that could be further investigated in future studies where more different weights should be measured.

More remarkably, we did find a similar object weight-specific effect of LO stimulation on load force rates: when stimulating LO in the dynamic phase, load force rates increased only for heavy objects. Such an effect was not seen in aIPS, where TMS over this area did not affect load force rates. This suggests that LO is specifically involved in the planning of load forces. In line with the narrative on weight discrimination, it seems that the load force planning also was less discriminative for object weight after LO stimulation. Correlations between stimulation effects on weight estimation and load force rates suggest that these effects could indeed be related. However, these correlations should be interpreted with care, since few correlations were seen and might not reflect stimulation effects but general similarities between load forces and weight estimation.

The effects on grip forces were not in line with effects of weight estimation or load force rates. Grip force rates increased after stimulation of either aIPS or LO, whereas static grip force was not significantly affected. Whereas previous research indicated that the amount of grip force applied during holding was related to weight estimation (Ellis and Lederman 1999; Flanagan and Bandomir 2000; Flanagan and Wing 1997), these might be governed by other areas than aIPS or LO as investigated in this study. While stimulation to both these areas increased grip force rates and reduced weight estimation, this effect was different regarding object weight and stimulation timing. The perceptual effect was only seen in heavy objects after stimulation in the static phase, whereas grip force rates were higher after stimulation in the dynamic phase both for light and heavy objects. It could be argued that the effect on grip force rates for both areas indicates a non-specific TMS effect rather than an actual indication of involvement of these areas (see limitations). However, the effect on grip forces we found was timing specific: only TMS applied in the dynamic loading phase increased grip forces, but not when applied in the static holding phase. In a previous study, a contribution of aIPS to grip force scaling was only found in a specific time window of 120-170 ms before object contact (Davare et al. 2007). The present results suggest that after object contact, both aIPS and LO play a role in the online control when grip forces increase, but not any longer when stable force levels are reached.

As mentioned before, whereas stimulation to both areas affected grip forces, load forces were only altered when TMS was applied over LO, not aIPS. This suggests that different processes are involved in planning of grip and load forces. In general, it has been assumed that grip and load forces are tightly coupled in an anticipatory manner (Flanagan and Wing 1993; Johansson and Westling 1988; 1984), suggesting they are controlled by the same neural network. Recently, however, it was shown that this coupling can be intermittent, not continuous (Grover et al. 2018). While both force components are important for object lifting, there are slight differences in their functionality. Grip forces are needed to ensure stability of the grasp to avoid slipping, which means they have to be adjusted to both object weight and surface friction and can contain a variable safety margin while still maintaining a stable grip. Load forces only need to be adjusted to object weight, which suggests a tighter link with weight perception, as indicated by our results. It must be noted that previous studies have found involvement of several brain areas for grip forces, but not load forces, such as left supplementary motor area (White et al. 2013), left aIPS (Davare et al. 2007) and dorsal premotor area (van Nuenen et al. 2012). To our knowledge, the neural correlates specifically tuned to load force scaling are less clear. Here we show for the first time an influence of a perceptual processing area, i.e. LO, on the control of load forces. Possibly, load and grip forces are generated with input from different brain areas and coupled together in others. In the literature, different areas have been suggested to be involved in this coupling, such as the cerebellum (Kawato et al. 2003), M1 (Schabrun et al. 2008) and the right intraparietal cortex (Ehrsson et al. 2003).

Finally, we hypothesized that the dynamic phase of object lifting would be more influential on weight perception than the static phase. Regarding force scaling, it has been shown that the lifting phase is important in building up sensorimotor memory (van Polanen and Davare 2019). Furthermore, weight perception is influenced by object size when the size is visually shown during lifting, but not holding and this effect reappears when the object is replaced (Plaisier et al. 2019). However, we did not find stronger effects of TMS on weight perception when it was applied in the dynamic phase compared to the static phase. Instead, significant effects were only seen for stimulation in the static phase for heavy objects. While this does not negate the importance of the dynamic phase for weight perception, it seems that LO and aIPS have a stronger influence on weight estimation in later phases of lifting.

The stimulation procedure as used in this study may also have some limitations. Because stimulation was provided randomly across trials, it is possible that TMS in the previous trial affected the storage of information differently for action and perceptual processes, thereby decreasing the relation between the two processes even on the following no-stimulation trials. However, we had too few trials to investigate the effects of stimulation in previous trials on the relation between force scaling and weight perception.

Furthermore, another limitation of the present study is that no sham stimulation condition was used. We used a control condition without stimulation to access normal baseline effects. Therefore, placebo effects of stimulation, such as auditory, somatosensory or startling effects from the TMS pulse, cannot be ruled out. Since we found effects of both aIPS and LO stimulation on grip forces, this could be due to a stimulation side effect. More specifically, participants could have squeezed more in response to the TMS and thereby increasing their grip force rates. We used two different TMS timings to control for such TMS side effects. Indeed, we only found an effect on grip forces when applying TMS in the dynamic, not the static phase. However, this does not completely rule out the possibility of a placebo effect in early phases of lifting because grip forces in dynamic and static phases might be differently susceptible to TMS. Further studies using appropriate sham conditions should rule out these possibilities.

Finally, since we only used two object weights in the study, the interpretation on differences between light and heavy objects is limited. Since the aim of the study was to investigate effects of previous lifts, using two weights was appropriate. However, to make definite conclusions about the effects of changes in weight discrimination and tuning of load forces to specific object weights, a larger range of object weights should be used.

To conclude, aIPS and LO do not seem to be involved in force scaling and weight judgement according to previous object experience. Whereas both areas appear to play a role in grip force scaling, LO specifically contributes to load force scaling, possibly related to object weight discrimination. This suggests that grip force and load forces are processed differently and both force components might be differently related to weight perception. More research is needed to shed more light on the relation between the involvement of LO in load force scaling and weight perception, and its potential role in weight discrimination.

## ACKNOWLEDGEMENTS

This research was supported by Fonds Wetenschappelijk Onderzoek grants (FWO postdoctoral fellowship, Belgium, 12X7118N; FWO Odysseus, Belgium, G/0C51/13N). The authors would like to thank members and students from the motor control and neuroplasticity lab for their help in data collection: L. Hermans, M. Gann, K. Heise, L. Pauwels, N. Dolfen and F. Monsieur.

## AUTHOR CONTRIBUTIONS

V.v.P. and M.D. conceived and designed research; V.v.P. and G.R. performed experiments; V.v.P. analyzed the data and prepared figures; V.v.P. and G.R. interpreted results of experiments; V.v.P. drafted manuscript; V.v.P., G.R. and M.D. edited and revised manuscript and approved final version of the manuscript.

