## Supplemental tables for "The role of the anterior intraparietal sulcus and the lateral occipital cortex in fingertip force scaling and weight perception during object lifting"

Supplementary Tables for Results and additional analysis in van Polanen, Rens, & Davare (2019). The role of the anterior intraparietal sulcus and the lateral occipital cortex in fingertip force scaling and weight perception during object lifting. bioRxiv 2019.12.20.883918; doi: https://doi.org/10.1101/2019.12.20.883918.

Table S1. ANOVA results for perceptual estimates, peak load force rate (peak LFR), time to peak LFR, first peak LFR (peak1 LFR), peak grip force rate (peak GFR), time to peak GFR, peak1 GFR and grip force during the static phase ($GF_{static}$). Factors are CW (current weight), PW (previous weight), TMS (TMS condition) and Loc (TMS location). The separate repeated measures ANOVAs were only performed when a significant effect or interaction with the between factor Loc was found. For the perceptual estimates this analysis was performed as an exploratory analysis for the trend (p=0.065) for the CW x TMS x Loc interaction. Significant effects are shown in blue.

### Perceptual estimates

| | Mixed ANOVA | | | ANOVA aIPS | | | ANOVA LO | | |
|---|---|---|---|---|---|---|---|---|---|
| **Factor** | **F (df, dfe)** | **p** | $\eta_p^2$ | **F (df, dfe)** | **p** | $\eta_p^2$ | **F (df, dfe)** | **p** | $\eta_p^2$ |
| CW | F(1,28)=66786.5 | <0.001 | 1 | F(1,14)=58349.6 | <0.001 | 1 | F(1,14)=23310.3 | <0.001 | 0.999 |
| PW | F(1,28)=0.5 | 0.002 | 0.294 | F(1,14)=6.9 | 0.02 | 0.332 | F(1,14)=4.8 | 0.046 | 0.256 |
| TMS | F(1,28)=11.7 | 0.008 | 0.183 | F(2,28)=5.0 | 0.014 | 0.263 | F(1,18)=2.2 | 0.15 | 0.136 |
| CW x PW | F(1,28)=0.1 | 0.197 | 0.059 | F(1,14)=1.3 | 0.275 | 0.085 | F(1,14)=0.5 | 0.48 | 0.036 |
| CW x TMS | F(1.5,41.9)=6.3 | 0.007 | 0.179 | F(2,28)=0.8 | 0.454 | 0.055 | F(2,28)=6.1 | 0.007 | 0.302 |
| PW x TMS | F(1,42)=0.4 | 0.207 | 0.055 | F(2,28)=0.6 | 0.574 | 0.039 | F(2,28)=1.1 | 0.348 | 0.073 |
| PW x CW x TMS | F(1,28)=1.7 | 0.632 | 0.016 | F(2,28)=2.1 | 0.138 | 0.132 | F(2,28)=0.0 | 0.98 | 0.001 |
| Loc | F(1,28)=0.8 | 0.379 | 0.028 | | | | | | |
| CW x Loc | F(1,28)=0.1 | 0.471 | 0.019 | | | | | | |
| PW x Loc | F(1.7,46.5)=6.1 | 0.741 | 0.004 | | | | | | |
| TMS x Loc | F(2,46)=3.1 | 0.633 | 0.013 | | | | | | |
| CW x PW x Loc | F(2,56)=1.6 | 0.758 | 0.003 | | | | | | |
| CW x TMS x Loc | F(2,56)=0.1 | 0.065 | 0.099 | | | | | | |
| PW x TMS x Loc | F(2,56)=0.5 | 0.863 | 0.005 | | | | | | |
| PW x CW x TMS x Loc | F(2,56)=0.8 | 0.44 | 0.029 | | | | | | |

### Peak LFR

| | Mixed ANOVA | | | ANOVA aIPS | | | ANOVA LO | | |
|---|---|---|---|---|---|---|---|---|---|
| **Factor** | **F (df, dfe)** | **p** | $\eta_p^2$ | **F (df, dfe)** | **p** | $\eta_p^2$ | **F (df, dfe)** | **p** | $\eta_p^2$ |
| CW | F(1,28)=171.8 | <0.001 | 0.86 | F(1,14)=63.2 | <0.001 | 0.819 | F(1,14)=140.0 | <0.001 | 0.909 |
| PW | F(1,28)=208.5 | <0.001 | 0.882 | F(1,14)=83.9 | <0.001 | 0.857 | F(1,14)=129.1 | <0.001 | 0.902 |
| TMS | F(1.3,35.2)=9.8 | 0.002 | 0.259 | F(1.3,17.9)=2.8 | 0.103 | 0.168 | F(1.2,16.7)=10.3 | 0.004 | 0.423 |
| CW x PW | F(1,28)=9.0 | 0.006 | 0.243 | F(1,14)=6.2 | 0.026 | 0.305 | F(1,14)=2.8 | 0.114 | 0.169 |
| CW x TMS | F(2,56)=4.9 | 0.01 | 0.15 | F(2,28)=0.2 | 0.79 | 0.017 | F(2,28)=9.6 | 0.001 | 0.407 |
| PW x TMS | F(2,56)=0.4 | 0.667 | 0.014 | F(2,28)=0.5 | 0.612 | 0.035 | F(2,28)=0.2 | 0.815 | 0.015 |
| PW x CW x TMS | F(2,56)=0.2 | 0.841 | 0.006 | F(2,28)=0.0 | 0.991 | 0.001 | F(2,28)=0.2 | 0.783 | 0.017 |
| Loc | F(1,28)=0.2 | 0.653 | 0.007 | | | | | | |
| CW x Loc | F(1,28)=0.1 | 0.816 | 0.002 | | | | | | |
| PW x Loc | F(1,28)=1.0 | 0.333 | 0.033 | | | | | | |
| TMS x Loc | F(1,35)=0.5 | 0.515 | 0.018 | | | | | | |
| CW x PW x Loc | F(1,28)=1.2 | 0.285 | 0.041 | | | | | | |
| CW x TMS x Loc | F(2,56)=3.2 | 0.048 | 0.103 | | | | | | |
| PW x TMS x Loc | F(2,56)=0.4 | 0.706 | 0.012 | | | | | | |
| PW x CW x TMS x Loc | F(2,56)=0.1 | 0.893 | 0.004 | | | | | | |

### Time to peak LFR

| | Mixed ANOVA | | | ANOVA aIPS | | | ANOVA LO | | |
|---|---|---|---|---|---|---|---|---|---|
| **Factor** | **F (df, dfe)** | **p** | $\eta_p^2$ | **F (df, dfe)** | **p** | $\eta_p^2$ | **F (df, dfe)** | **p** | $\eta_p^2$ |
| CW | F(1,28)=246.7 | <0.001 | 0.898 | F(1,14)=115.0 | <0.001 | 0.891 | F(1,14)=136.6 | <0.001 | 0.907 |
| PW | F(1,28)=0.1 | 0.728 | 0.004 | F(1,14)=0.0 | 0.942 | 0 | F(1,14)=0.2 | 0.663 | 0.014 |
| TMS | F(2,56)=3.5 | 0.037 | 0.111 | F(2,28)=0.5 | 0.611 | 0.035 | F(2,28)=4.1 | 0.028 | 0.225 |
| CW x PW | F(1,28)=4.9 | 0.035 | 0.149 | F(1,14)=3.6 | 0.079 | 0.204 | F(1,14)=1.3 | 0.267 | 0.087 |
| CW x TMS | F(1.6,45.7)=4.9 | 0.016 | 0.15 | F(2,28)=0.5 | 0.621 | 0.034 | F(2,28)=6.9 | 0.004 | 0.332 |
| PW x TMS | F(2,56)=1.1 | 0.331 | 0.039 | F(2,28)=0.7 | 0.527 | 0.045 | F(2,28)=0.7 | 0.498 | 0.049 |
| PW x CW x TMS | F(2,56)=0.9 | 0.414 | 0.031 | F(2,28)=0.0 | 0.952 | 0.003 | F(2,28)=1.3 | 0.298 | 0.083 |
| Loc | F(1,28)=8.7 | 0.006 | 0.236 | | | | | | |
| CW x Loc | F(1,28)=0.7 | 0.4 | 0.025 | | | | | | |
| PW x Loc | F(1,28)=0.1 | 0.812 | 0.002 | | | | | | |
| TMS x Loc | F(2,56)=0.7 | 0.519 | 0.023 | | | | | | |
| CW x PW x Loc | F(1,28)=1.1 | 0.295 | 0.039 | | | | | | |
| CW x TMS x Loc | F(2,46)=1.8 | 0.182 | 0.06 | | | | | | |
| PW x TMS x Loc | F(2,56)=0.2 | 0.789 | 0.008 | | | | | | |
| PW x CW x TMS x Loc | F(2,56)=0.5 | 0.628 | 0.017 | | | | | | |

### Peak1 LFR

| | Mixed ANOVA | | | ANOVA aIPS | | | ANOVA LO | | |
|---|---|---|---|---|---|---|---|---|---|
| **Factor** | **F (df, dfe)** | **p** | $\eta_p^2$ | **F (df, dfe)** | **p** | $\eta_p^2$ | **F (df, dfe)** | **p** | $\eta_p^2$ |
| CW | F(1,28)=53.9 | <0.001 | 0.658 | | | | | | |
| PW | F(1,28)=137.1 | <0.001 | 0.83 | | | | | | |
| TMS | F(1.6,43.8)=1.7 | 0.202 | 0.057 | | | | | | |
| CW x PW | F(1,28)=4.8 | 0.036 | 0.147 | | | | | | |
| CW x TMS | F(2,56)=0.6 | 0.578 | 0.019 | | | | | | |
| PW x TMS | F(2,56)=1.2 | 0.314 | 0.041 | | | | | | |
| PW x CW x TMS | F(2,56)=0.1 | 0.882 | 0.004 | | | | | | |
| Loc | F(1,28)=0.4 | 0.526 | 0.015 | | | | | | |
| CW x Loc | F(1,28)=0.0 | 0.906 | 0.001 | | | | | | |
| PW x Loc | F(1,28)=0.9 | 0.343 | 0.032 | | | | | | |
| TMS x Loc | F(2,44)=0.6 | 0.531 | 0.02 | | | | | | |
| CW x PW x Loc | F(1,28)=0.3 | 0.583 | 0.011 | | | | | | |
| CW x TMS x Loc | F(2,56)=1.8 | 0.182 | 0.059 | | | | | | |
| PW x TMS x Loc | F(2,56)=0.3 | 0.767 | 0.009 | | | | | | |
| PW x CW x TMS x Loc | F(2,56)=0.0 | 0.999 | 0 | | | | | | |

### Peak GFR

| | Mixed ANOVA | | | ANOVA aIPS | | | ANOVA LO | | |
|---|---|---|---|---|---|---|---|---|---|
| **Factor** | **F (df, dfe)** | **p** | $\eta_p^2$ | **F (df, dfe)** | **p** | $\eta_p^2$ | **F (df, dfe)** | **p** | $\eta_p^2$ |
| CW | F(1,28)=44.6 | <0.001 | 0.614 | F(1,14)=43.0 | <0.001 | 0.754 | F(1,14)=14.4 | 0.002 | 0.506 |
| PW | F(1,28)=204.9 | <0.001 | 0.88 | F(1,14)=54.1 | <0.001 | 0.795 | F(1,14)=228.6 | <0.001 | 0.942 |
| TMS | F(1.5,41.1)=18.4 | <0.001 | 0.396 | F(1.3,18.9)=8.0 | 0.007 | 0.365 | F(2,28)=14.7 | <0.001 | 0.513 |
| CW x PW | F(1,28)=0.0 | 0.878 | 0.001 | F(1,14)=0.3 | 0.566 | 0.024 | F(1,14)=0.6 | 0.466 | 0.039 |
| CW x TMS | F(2,56)=3.6 | 0.035 | 0.113 | F(2,28)=0.9 | 0.429 | 0.059 | F(2,28)=3.0 | 0.064 | 0.179 |
| PW x TMS | F(1.6,44.6)=2.9 | 0.076 | 0.094 | F(2,28)=1.6 | 0.222 | 0.102 | F(2,28)=1.5 | 0.247 | 0.095 |
| PW x CW x TMS | F(2,56)=0.1 | 0.872 | 0.005 | F(2,28)=0.5 | 0.621 | 0.033 | F(2,28)=1.0 | 0.385 | 0.066 |
| Loc | F(1,28)=4.4 | 0.046 | 0.135 | | | | | | |
| CW x Loc | F(1,28)=0.1 | 0.813 | 0.002 | | | | | | |
| PW x Loc | F(1,28)=3.6 | 0.069 | 0.113 | | | | | | |
| TMS x Loc | F(1,41)=0.7 | 0.475 | 0.023 | | | | | | |
| CW x PW x Loc | F(1,28)=0.9 | 0.351 | 0.031 | | | | | | |
| CW x TMS x Loc | F(2,56)=0.3 | 0.715 | 0.012 | | | | | | |
| PW x TMS x Loc | F(2,45)=0.2 | 0.8 | 0.006 | | | | | | |
| PW x CW x TMS x Loc | F(2,56)=1.4 | 0.243 | 0.049 | | | | | | |

### Time to peak GFR

| | Mixed ANOVA | | | ANOVA aIPS | | | ANOVA LO | | |
|---|---|---|---|---|---|---|---|---|---|
| **Factor** | **F (df, dfe)** | **p** | $\eta_p^2$ | **F (df, dfe)** | **p** | $\eta_p^2$ | **F (df, dfe)** | **p** | $\eta_p^2$ |
| CW | F(1,28)=60.0 | <0.001 | 0.682 | | | | | | |
| PW | F(1,28)=2.8 | 0.105 | 0.091 | | | | | | |
| TMS | F(2,56)=1.5 | 0.231 | 0.051 | | | | | | |
| CW x PW | F(1,28)=16.4 | <0.001 | 0.37 | | | | | | |
| CW x TMS | F(2,56)=1.3 | 0.284 | 0.044 | | | | | | |
| PW x TMS | F(2,56)=0.5 | 0.618 | 0.017 | | | | | | |
| PW x CW x TMS | F(2,56)=0.5 | 0.635 | 0.016 | | | | | | |
| Loc | F(1,28)=2.9 | 0.101 | 0.093 | | | | | | |
| CW x Loc | F(1,28)=0.0 | 0.945 | 0 | | | | | | |
| PW x Loc | F(1,28)=0.1 | 0.738 | 0.004 | | | | | | |
| TMS x Loc | F(2,56)=1.1 | 0.331 | 0.039 | | | | | | |
| CW x PW x Loc | F(1,28)=0.6 | 0.448 | 0.021 | | | | | | |
| CW x TMS x Loc | F(2,56)=1.0 | 0.381 | 0.034 | | | | | | |
| PW x TMS x Loc | F(2,56)=0.1 | 0.934 | 0.002 | | | | | | |
| PW x CW x TMS x Loc | F(2,56)=1.1 | 0.353 | 0.037 | | | | | | |

### Peak1 GFR

| | Mixed ANOVA | | | ANOVA aIPS | | | ANOVA LO | | |
|---|---|---|---|---|---|---|---|---|---|
| **Factor** | **F (df, dfe)** | **p** | $\eta_p^2$ | **F (df, dfe)** | **p** | $\eta_p^2$ | **F (df, dfe)** | **p** | $\eta_p^2$ |
| CW | F(1,28)=13.4 | 0.001 | 0.324 | F(1,14)=19.2 | 0.001 | 0.579 | F(1,14)=2.1 | 0.166 | 0.133 |
| PW | F(1,28)=170.7 | <0.001 | 0.859 | F(1,14)=43.3 | <0.001 | 0.756 | F(1,14)=187.9 | <0.001 | 0.931 |
| TMS | F(2,56)=6.5 | 0.003 | 0.189 | F(2,28)=5.0 | 0.014 | 0.263 | F(2,28)=1.7 | 0.203 | 0.108 |
| CW x PW | F(1,28)=2.3 | 0.14 | 0.076 | F(1,14)=0.3 | 0.568 | 0.024 | F(1,14)=2.3 | 0.154 | 0.14 |
| CW x TMS | F(2,56)=0.2 | 0.82 | 0.007 | F(2,28)=0.2 | 0.835 | 0.013 | F(2,28)=0.4 | 0.667 | 0.029 |
| PW x TMS | F(1.6,45.7)=3.2 | 0.058 | 0.103 | F(2,28)=1.2 | 0.313 | 0.08 | F(2,28)=2.1 | 0.142 | 0.13 |
| PW x CW x TMS | F(2,56)=0.5 | 0.581 | 0.019 | F(2,28)=1.5 | 0.243 | 0.096 | F(2,28)=1.1 | 0.332 | 0.076 |
| Loc | F(1,28)=4.2 | 0.050 | 0.13 | | | | | | |
| CW x Loc | F(1,28)=1.5 | 0.229 | 0.051 | | | | | | |
| PW x Loc | F(1,28)=4.4 | 0.045 | 0.136 | | | | | | |
| TMS x Loc | F(2,56)=2.0 | 0.146 | 0.066 | | | | | | |
| CW x PW x Loc | F(1,28)=0.6 | 0.462 | 0.019 | | | | | | |
| CW x TMS x Loc | F(2,56)=0.4 | 0.684 | 0.013 | | | | | | |
| PW x TMS x Loc | F(2,46)=0.1 | 0.909 | 0.002 | | | | | | |
| PW x CW x TMS x Loc | F(2,56)=2.0 | 0.151 | 0.065 | | | | | | |

### $GF_{static}$

| | Mixed ANOVA | | | ANOVA aIPS | | | ANOVA LO | | |
|---|---|---|---|---|---|---|---|---|---|
| **Factor** | **F (df, dfe)** | **p** | $\eta_p^2$ | **F (df, dfe)** | **p** | $\eta_p^2$ | **F (df, dfe)** | **p** | $\eta_p^2$ |
| CW | F(1,28)=1276.0 | <0.001 | 0.979 | F(1,14)=757.4 | <0.001 | 0.982 | F(1,14)=570.6 | <0.001 | 0.976 |
| PW | F(1,28)=12.7 | 0.001 | 0.312 | F(1,14)=1.4 | 0.264 | 0.088 | F(1,14)=31.8 | <0.001 | 0.695 |
| TMS | F(2,56)=2.4 | 0.096 | 0.08 | F(2,28)=1.3 | 0.284 | 0.086 | F(2,28)=1.1 | 0.337 | 0.075 |
| CW x PW | F(1,28)=2.8 | 0.106 | 0.091 | F(1,14)=0.1 | 0.724 | 0.009 | F(1,14)=15.7 | 0.001 | 0.529 |
| CW x TMS | F(2,56)=7.5 | 0.001 | 0.211 | F(1.4,19.5)=4.6 | 0.034 | 0.246 | F(2,28)=3.3 | 0.051 | 0.191 |
| PW x TMS | F(2,56)=1.2 | 0.32 | 0.04 | F(2,28)=0.0 | 0.966 | 0.002 | F(2,28)=2.2 | 0.124 | 0.138 |
| PW x CW x TMS | F(2,56)=0.1 | 0.87 | 0.005 | F(1.4,19.5)=0.7 | 0.444 | 0.05 | F(2,28)=0.3 | 0.772 | 0.018 |
| Loc | F(1,28)=8.8 | 0.006 | 0.238 | | | | | | |
| CW x Loc | F(1,28)=4.5 | 0.044 | 0.138 | | | | | | |
| PW x Loc | F(1,28)=2.2 | 0.151 | 0.072 | | | | | | |
| TMS x Loc | F(2,56)=0.0 | 0.981 | 0.001 | | | | | | |
| CW x PW x Loc | F(1,28)=5.3 | 0.029 | 0.158 | | | | | | |
| CW x TMS x Loc | F(2,56)=0.1 | 0.877 | 0.005 | | | | | | |
| PW x TMS x Loc | F(2,56)=1.3 | 0.281 | 0.044 | | | | | | |
| PW x CW x TMS x Loc | F(2,56)=0.9 | 0.415 | 0.031 | | | | | | |

Table S2. ANOVA results for the effects of TMS in the previous trial on the current trial. Results are shown for perceptual estimates, peak load force rate (peak LFR), peak grip force rate (peak GFR) and grip force during the static phase ($GF_{static}$). Factors are CW (current weight), PW (previous weight), TMS (TMS condition in previous trial) and Loc (TMS location in previous trial). The separate repeated measures ANOVAs were only performed when a significant effect or interaction with the between factor Loc was found. Significant effects are shown in blue.

### Perceptual estimates

| | Mixed ANOVA | | | ANOVA aIPS | | | ANOVA LO | | |
|---|---|---|---|---|---|---|---|---|---|
| **Factor** | **F (df, dfe)** | **p** | $\eta_p^2$ | **F (df, dfe)** | **p** | $\eta_p^2$ | **F (df, dfe)** | **p** | $\eta_p^2$ |
| CW | F(1,28)=54825.5 | <0.001 | 0.999 | | | | | | |
| PW | F(1,28)=10.4 | 0.003 | 0.271 | | | | | | |
| TMS | F(2,56)=2.3 | 0.113 | 0.075 | | | | | | |
| CW x PW | F(1,28)=2.2 | 0.146 | 0.074 | | | | | | |
| CW x TMS | F(1.6,44.2)=1.6 | 0.216 | 0.054 | | | | | | |
| PW x TMS | F(1.5,42.4)=0.3 | 0.712 | 0.009 | | | | | | |
| PW x CW x TMS | F(2,56)=0.5 | 0.639 | 0.016 | | | | | | |
| Loc | F(1,28)=1.2 | 0.287 | 0.04 | | | | | | |
| CW x Loc | F(1,28)=0.1 | 0.716 | 0.005 | | | | | | |
| PW x Loc | F(1,28)=0.2 | 0.649 | 0.008 | | | | | | |
| TMS x Loc | F(2,56)=0.5 | 0.586 | 0.019 | | | | | | |
| CW x PW x Loc | F(1,28)=0.1 | 0.702 | 0.005 | | | | | | |
| CW x TMS x Loc | F(1.6,44.2)=0.3 | 0.724 | 0.009 | | | | | | |
| PW x TMS x Loc | F(1.5,42.4)=0.9 | 0.388 | 0.031 | | | | | | |
| PW x CW x TMS x Loc | F(2,56)=0.2 | 0.856 | 0.006 | | | | | | |

### Peak LFR

| | Mixed ANOVA | | | ANOVA aIPS | | | ANOVA LO | | |
|---|---|---|---|---|---|---|---|---|---|
| **Factor** | **F (df, dfe)** | **p** | $\eta_p^2$ | **F (df, dfe)** | **p** | $\eta_p^2$ | **F (df, dfe)** | **p** | $\eta_p^2$ |
| CW | F(1,28)=153.6 | <0.001 | 0.846 | | | | | | |
| PW | F(1,28)=217.2 | <0.001 | 0.886 | | | | | | |
| TMS | F(1.5,43.1)=0.2 | 0.796 | 0.006 | | | | | | |
| CW x PW | F(1,28)=9.3 | 0.005 | 0.249 | | | | | | |
| CW x TMS | F(2,56)=0.1 | 0.945 | 0.002 | | | | | | |
| PW x TMS | F(2,56)=1.7 | 0.19 | 0.058 | | | | | | |
| PW x CW x TMS | F(2,56)=0.5 | 0.622 | 0.017 | | | | | | |
| Loc | F(1,28)=0.2 | 0.677 | 0.006 | | | | | | |
| CW x Loc | F(1,28)=0.1 | 0.767 | 0.003 | | | | | | |
| PW x Loc | F(1,28)=1.0 | 0.32 | 0.035 | | | | | | |
| TMS x Loc | F(1.5,43.1)=1.0 | 0.365 | 0.034 | | | | | | |
| CW x PW x Loc | F(1,28)=1.4 | 0.251 | 0.047 | | | | | | |
| CW x TMS x Loc | F(2,56)=0.1 | 0.873 | 0.005 | | | | | | |
| PW x TMS x Loc | F(2,56)=0.8 | 0.435 | 0.029 | | | | | | |
| PW x CW x TMS x Loc | F(2,56)=2.0 | 0.145 | 0.067 | | | | | | |

### Peak GFR

| | Mixed ANOVA | | | ANOVA aIPS | | | ANOVA LO | | |
|---|---|---|---|---|---|---|---|---|---|
| **Factor** | **F (df, dfe)** | **p** | $\eta_p^2$ | **F (df, dfe)** | **p** | $\eta_p^2$ | **F (df, dfe)** | **p** | $\eta_p^2$ |
| CW | F(1,28)=37.2 | <0.001 | 0.57 | F(1,14)=35.0 | <0.001 | 0.714 | F(1,14)=11.7 | 0.004 | 0.456 |
| PW | F(1,28)=213.7 | <0.001 | 0.884 | F(1,14)=58.8 | <0.001 | 0.808 | F(1,14)=225.5 | <0.001 | 0.942 |
| TMS | F(2,56)=2.4 | 0.101 | 0.079 | F(2,28)=3.2 | 0.054 | 0.188 | F(2,28)=0.4 | 0.698 | 0.025 |
| CW x PW | F(1,28)=0.2 | 0.661 | 0.007 | F(1,14)=0.3 | 0.565 | 0.024 | F(1,14)=1.4 | 0.251 | 0.093 |
| CW x TMS | F(1.6,45.6)=0.4 | 0.603 | 0.016 | F(2,28)=0.5 | 0.61 | 0.035 | F(2,28)=3.0 | 0.065 | 0.177 |
| PW x TMS | F(2,56)=4.2 | 0.02 | 0.131 | F(2,28)=0.7 | 0.503 | 0.048 | F(2,28)=6.0 | 0.007 | 0.301 |
| PW x CW x TMS | F(2,56)=1.1 | 0.347 | 0.037 | F(2,28)=5.5 | 0.01 | 0.28 | F(2,28)=0.6 | 0.563 | 0.04 |
| Loc | F(1,28)=4.4 | 0.046 | 0.135 | | | | | | |
| CW x Loc | F(1,28)=0.1 | 0.725 | 0.004 | | | | | | |
| PW x Loc | F(1,28)=3.1 | 0.089 | 0.1 | | | | | | |
| TMS x Loc | F(2,56)=0.8 | 0.465 | 0.027 | | | | | | |
| CW x PW x Loc | F(1,28)=1.6 | 0.215 | 0.054 | | | | | | |
| CW x TMS x Loc | F(1.6,45.6)=2.2 | 0.13 | 0.073 | | | | | | |
| PW x TMS x Loc | F(2,56)=2.8 | 0.072 | 0.09 | | | | | | |
| PW x CW x TMS x Loc | F(2,56)=2.7 | 0.079 | 0.087 | | | | | | |

### $GF_{static}$

| | Mixed ANOVA | | | ANOVA aIPS | | | ANOVA LO | | |
|---|---|---|---|---|---|---|---|---|---|
| **Factor** | **F (df, dfe)** | **p** | $\eta_p^2$ | **F (df, dfe)** | **p** | $\eta_p^2$ | **F (df, dfe)** | **p** | $\eta_p^2$ |
| CW | F(1,28)=1272.1 | <0.001 | 0.978 | F(1,14)=736.4 | <0.001 | 0.981 | F(1,14)=577.0 | <0.001 | 0.976 |
| PW | F(1,28)=12.5 | 0.001 | 0.308 | F(1,14)=1.5 | 0.241 | 0.097 | F(1,14)=28.2 | <0.001 | 0.668 |
| TMS | F(2,56)=3.7 | 0.032 | 0.116 | F(2,28)=1.8 | 0.177 | 0.116 | F(1.4,20.1)=2.2 | 0.149 | 0.135 |
| CW x PW | F(1,28)=2.6 | 0.119 | 0.084 | F(1,14)=0.1 | 0.741 | 0.008 | F(1,14)=11.0 | 0.005 | 0.441 |
| CW x TMS | F(2,56)=0.5 | 0.611 | 0.017 | F(2,28)=0.4 | 0.702 | 0.025 | F(2,28)=0.8 | 0.481 | 0.051 |
| PW x TMS | F(1.6,45.2)=2.6 | 0.095 | 0.085 | F(2,28)=3.3 | 0.053 | 0.189 | F(1.4,20.2)=0.5 | 0.546 | 0.036 |
| PW x CW x TMS | F(2,56)=0.4 | 0.652 | 0.015 | F(2,28)=0.1 | 0.918 | 0.006 | F(2,28)=0.8 | 0.481 | 0.051 |
| Loc | F(1,28)=8.7 | 0.006 | 0.238 | | | | | | |
| CW x Loc | F(1,28)=4.2 | 0.05 | 0.131 | | | | | | |
| PW x Loc | F(1,28)=1.8 | 0.186 | 0.062 | | | | | | |
| TMS x Loc | F(2,56)=0.4 | 0.691 | 0.013 | | | | | | |
| CW x PW x Loc | F(1,28)=4.7 | 0.039 | 0.143 | | | | | | |
| CW x TMS x Loc | F(2,56)=0.5 | 0.631 | 0.016 | | | | | | |
| PW x TMS x Loc | F(1.6,45.2)=0.4 | 0.617 | 0.015 | | | | | | |
| PW x CW x TMS x Loc | F(2,56)=0.8 | 0.475 | 0.026 | | | | | | |

Table S3. Results for the mixed linear model to investigate the relation between force rates and perceptual estimates. Factors are Loc (TMS location), TMS (TMS condition), CW (current weight) and FR (peak force rate). F-values, with degrees of freedom for the numerator and denominator, and p-values are shown. Significant effects are shown in blue.

| | Previous lift | | TMS | | Trial-by-trial | |
|---|---|---|---|---|---|---|
| | Peak GFR | LFRmax | GFRmax | LFRmax | GFRmax | LFRmax |
| Intercept | F(1,33)=4.8, p=0.035 | F(1,36)=2.6, p=0.115 | F(1,19)=6.6, p=0.019 | F(1,15)=7.0, p=0.018 | F(1,15)=1.6, p=0.222 | F(1,16)=1.1, p=0.307 |
| Loc | F(1,174)=1.9, p=0.173 | F(1,106)=0.3, p=0.582 | F(1,107)=0.1, p=0.747 | F(1,103)=1.6, p=0.211 | F(1,4004)=2.5, p=0.118 | F(1,4005)=2.9, p=0.088 |
| TMS | F(1,127)=3.4, p=0.067 | F(1,141)=1.8, p=0.181 | F(1,106)=17.4, p<0.001 | F(1,102)=11.4, p=0.001 | F(2,3416)=10.4, p<0.001 | F(2,3484)=11.0, p<0.001 |
| CW | F(2,142)=0.5, p=0.634 | F(2,137)=1.0, p=0.387 | F(1,108)=0.1, p=0.805 | F(1,102)<0.1, p=0.968 | F(1,3415)=63290, p<0.001 | F(1,3488)=59074, p<0.001 |
| FR | F(1,152)=0.1, p=0.714 | F(1,164)=2.6, p=0.109 | F(1,119)=1.3, p=0.258 | F(1,115)=3.6, p=0.061 | F(1,3254)=11.7, p=0.001 | F(1,3631)=7.0, p=0.008 |
| Loc x TMS | F(1,167)<0.1, p=0.948 | F(1,107)=0.2, p=0.656 | F(1,106)=4.6, p=0.034 | F(1,103)=2.0, p=0.160 | F(2,4001)=0.7, p=0.492 | F(2,4005)=0.5, p=0.610 |
| Loc x CW | F(2,167)=0.6, p=0.554 | F(2,104)=0.6, p=0.572 | F(1,108)=1.3, p=0.251 | F(1,102)<0.1, p=0.837 | F(1,4004)=0.8, p=0.369 | F(1,4005)<0.1, p=0.916 |
| loc x FR | F(1,174)=1.3, p=0.257 | F(1,149)=0.1, p=0.722 | F(1,114)<0.1, p=0.997 | F(1,111)=5.0, p=0.027 | F(1,4007)<0.1, p=0.836 | F(1,4006)=2.8, p=0.094 |
| TMS x CW | F(2,138)=1.1, p=0.323 | F(2,140)=0.3, p=0.708 | F(1,104)=0.3, p=0.586 | F(1,108)<0.1, p=0.871 | F(2,3417)=8.4, p<0.001 | F(2,3484)=6.8, p=0.001 |
| TMS x FR | F(1,146)=1.0, p=0.324 | F(1,158)=0.1, p=0.817 | F(1,113)<0.1, p=0.896 | F(1,113)=0.2, p=0.647 | F(2,3267)=0.4, p=0.674 | F(2,3596)=1.2, p=0.300 |
| CW x FR | F(2,158)=0.3, p=0.769 | F(2,149)=0.5, p=0.580 | F(1,108)=0.7, p=0.407 | F(1,109)=1.0, p=0.308 | F(1,3266)=1.3, p=0.246 | F(1,3586)<0.1, p=0.972 |
| Loc x TMS x CW | F(2,169)=0.7, p=0.476 | F(2,103)=0.8, p=0.465 | F(1,105)=0.3, p=0.614 | F(1,99)=0.1, p=0.771 | F(2,4002)=4.1, p=0.017 | F(2,4005)=3.3, p=0.039 |
| Loc x TMS x FR | F(1,168)=0.1, p=0.787 | F(1,145)=0.1, p=0.761 | F(1,110)=3.4, p=0.068 | F(1,110)=7.3, p=0.008 | F(2,4014)=0.2, p=0.811 | F(2,4004)=0.2, p=0.819 |
| Loc x CW x FR | F(2,167)=0.4, p=0.657 | F(2,145)=0.2, p=0.826 | F(1,112)=0.8, p=0.364 | F(1,109)=0.6, p=0.422 | F(1,4016)<0.1, p=0.961 | F(1,4006)=0.1, p=0.737 |
| TMS x CW x FR | F(2,159)=1.9, p=0.149 | F(2,156)=0.1, p=0.923 | F(1,109)=0.9, p=0.356 | F(1,118)=2.8, p=0.100 | F(2,3252)=0.6, p=0.563 | F(2,3621)=0.2, p=0.800 |
| Loc x TMS x CW x FR | F(2,170)=0.6, p=0.557 | F(2,146)=0.9, p=0.400 | F(1,106)=2.2, p=0.141 | F(1,109)=0.3, p=0.561 | F(2,4006)=1.0, p=0.376 | F(2,4004)=0.2, p=0.852 |
